# Adaptation to recent outcomes attenuates the lasting effect of initial experience on risky decisions

**DOI:** 10.1101/2020.09.25.313551

**Authors:** Andrea Kóbor, Zsófia Kardos, Ádám Takács, Noémi Éltető, Karolina Janacsek, Eszter Tóth-Fáber, Valéria Csépe, Dezso Nemeth

## Abstract

Both primarily and recently encountered information have been shown to influence experience-based risky decision making. The primacy effect predicts that initial experience will influence later choices even if outcome probabilities change and reward is ultimately more or less sparse than primarily experienced. However, it has not been investigated whether extended initial experience would induce a more profound primacy effect upon risky choices than brief experience. Therefore, the present study tested in two experiments whether young adults adjusted their risk-taking behavior in the Balloon Analogue Risk task after an unsignaled and unexpected change point. The change point separated early “good luck” or “bad luck” trials from subsequent ones. While mostly positive (more reward) or mostly negative (no reward) events characterized the early trials, subsequent trials were unbiased. In Experiment 1, the change point occurred after one-sixth or one-third of the trials (brief vs. extended experience) without intermittence, whereas in Experiment 2, it occurred between separate task phases. In Experiment 1, if negative events characterized the early trials, after the change point, risk-taking behavior increased as compared with the early trials. Conversely, if positive events characterized the early trials, risk-taking behavior decreased after the change point. Although the adjustment of risk-taking behavior occurred due to integrating recent experiences, the impact of initial experience was simultaneously observed. The length of initial experience did not reliably influence the adjustment of behavior. In Experiment 2, participants became more prone to take risks as the task progressed, indicating that the impact of initial experience could be overcome. Altogether, we suggest that initial beliefs about outcome probabilities can be updated by recent experiences to adapt to the continuously changing decision environment.

## Introduction

Decision making is supported by generating predictions about the probability of upcoming events. Many of these predictions are guided by inferences based on prior experiences with a similar situation or early experiences with the current situation. Considering risky choices, if we learned that regulations were moderate and had the repeated experience that driving beyond the speed limit did not end in negative consequences, we would be more likely to continue this behavior. However, in the case of stricter regulations, this habitual choice behavior becomes disadvantageous and should be overcome. Adaptation to the newly encountered regulations is probably gradual but takes an unknown amount of time, depending on several factors. Accordingly, this study employs an experience-based risky decision-making paradigm to investigate whether initial experience or the combination of initial and recent experiences had stronger impact on risk-taking behavior after an unsignaled change in outcome probabilities.

When decisions are made under uncertainty, such as in the present study, the probability distribution of the possible outcomes is unknown ^1–5^. The probability distribution can be approximated via repeated sampling and inference; therefore, these decisions are based on experience instead of description ^6, 7^. Decisions made under uncertainty differ from those made under explicit risk. In the latter case, although outcomes are unknown, outcome probabilities are described a priori and known by the individuals. Consequently, these two types of decisions differ in their underlying processes, as well. For instance, decisions under uncertainty are less dependent on executive functions and involve the use of heuristics or somatic markers (for review, see ^1, 4^). In day-to-day life, we make decisions mostly from experience in uncertain situations.

It has long been investigated how and under which conditions belief updating about outcome probabilities occurs in various decision-making situations (cf. ^8^). For instance, the temporal order of information seems to play a considerable role when decisions are uncertain and based on experience ^6^. According to the anchoring and adjustment heuristic, initial beliefs based on limited observations serve as a reference point that is adjusted as information accumulates. The initial anchor’s effect is probably long-lasting since adjustments are usually insufficient and biased toward the initial values ^9^. Meanwhile, the complexity of the task, the amount of new evidence, and the response mode (responding after integrating each piece of evidence or after the presentation of all pieces of evidence) have been shown to influence the effect of information order ^8^. Thus, there are conditions under which primacy, recency, or no order effects, respectively, are more likely to occur.

Evidence from studies targeting or modeling real-life risky situations supports the combination of both primacy and recency effects. Based on survey data, it was shown that those who had experienced great macroeconomic shocks became financially more risk averse in their later lives; for instance, they were less likely to participate in the stock market ^10^. Although recent experiences received more weight than more distant ones, the impact of experiences that had happened even years earlier remained significant. This so-called depression baby hypothesis has also been tested with simulated experimental stock market experiments where participants hypothetically invested in safe (cash deposit account) and risky (stock index fund) options ^11^. These experiments manipulated whether participants learned from experience (played the investment game) or from symbolic description (only studied graphs of the initial period) and whether the data included or excluded initial macroeconomic crisis (negative shock) or boom (positive shock). Depending on the valence of the initial shock, participants who learned from experience took less or more risks than those who learned from description (cf. ^6, 12^). Importantly, the most recent experience also played a role in participants’ investment behavior that was reactive to local fluctuations in stock prices, irrespective of experimental manipulations.

Other studies of sequential decision making indicate the prominence of recency effect ^13^. Due to the continuous tracking of the recent outcomes, in nonstationary stochastic environments, individuals accurately estimated the underlying probabilities and quickly updated their estimates if there was a change in these probabilities ^14–16^. The prominent impact of recently observed outcomes was also demonstrated for the sequential sampling of gambles and their valuation ^17, 18^.

The studies described so far suggest that individuals could successfully use recently observed evidence to adapt to the changing environment (cf. ^19^). However, fast adaptation to initial payoff probabilities followed by only slow adaptation to changes in these probabilities was also found, especially if the history summarizing previous outcomes was explicitly provided to participants ^20, 21^. In addition, as demonstrated in a hypothetical stock market game, the initially established decision strategy was retained even though it became no longer optimal ^22^. A hint about the change, monetary incentives, or transfer to another task with the same deep structure but different surface could not adequately support the update of this original decision strategy. Therefore, depending on the decision situation, belief updating could also be difficult or completely missing.

Among tasks requiring sequential decisions, the Balloon Analogue Risk Task (BART) has been a widely used laboratory measure because its structure and appearance mimics naturalistic risk taking ^23–26^. This task requires sequential decisions on risk in the form of inflating virtual balloons on a screen. Participants are told that each balloon pump is associated with either a balloon inflation or a balloon burst. While balloon inflations are followed by reward (increasing score), balloon bursts are followed by no reward (loss of score accumulated on the given balloon). However, the outcome of each successive pump (inflation vs. burst) and the probability distribution determining the outcome are unknown to participants. As the task progresses, participants could learn about the payoff structure in a trial-and-error manner to maximize the total reward. Thus, the BART measures decision making under uncertainty and experience-based risk, at least during the early trials ^1–4, 27^.

To our knowledge, two studies investigated whether beliefs about outcome probabilities are updated if these probabilities change considerably after a certain balloon in the BART. The study of Koscielniak, et al. ^28^ showed that initial good luck and bad luck modulated subsequent adaptation. After the unlucky series (bursts on the first three balloons after three, four, and one pumps, irrespective of the participant’s choice), the number of balloon inflations gradually increased in the following phases of the task. After the lucky series (bursts on the first three balloons after 25, 27, and 28 pumps), their number was higher only in the first phase following manipulation and decreased in the middle and final phases. However, the number of balloon inflations in the bad luck and good luck conditions remained different in every phase. The study of Bonini, et al. ^29^ used five early losses (bursts at the first, second, third, second, and first pumps) in a modified BART and contrasted the observed balloon inflations with that of an original BART. Healthy control participants showed a lower number of balloon inflations across the remaining balloons of the modified BART as compared with the original BART. As the task progressed, they increased the number of balloon inflations, and this increase was steeper in the modified BART.

In sum, although early negative events in the BART decreased overall risk taking (i.e., fewer balloon inflations), experience with more balloons still increased it gradually ^10, 11^. Nevertheless, previous studies manipulated only the very first trials by using extremely high or low balloon tolerances. Therefore, it has not been systematically tested whether longer initial experience might result in more persistent primacy effect and weaker subsequent adaptation to the changed payoff structure than shorter experience, in line with the existing accounts (cf. ^8, 10, 20^).

Accordingly, the present study investigated how different lengths of initial experience influenced risk-taking behavior during sequential sampling of outcome probabilities without signaling the change in the payoff structure. In Experiment 1, the first five (short manipulation) or ten (long manipulation) balloons of the task consisting of 30 balloons were manipulated to induce negative or positive initial experience. While the results of Experiment 1 suggested that different lengths of both negative and positive initial experience could be overcome to a limited extent, two questions remained unanswered. First, because the underlying payoff structure changed at different times across the long and short experimental conditions, participants encountered a slightly different series of unbiased balloons after the manipulated phase. Therefore, the potential to change risk taking on the unbiased balloons was differently constrained across the length conditions. Second, without baseline experience (i.e., a reference level), whether lucky and unlucky experiences indeed led to more and less risk taking was not measured directly (cf. ^30^). To address these issues, in Experiment 2, the manipulated phase was separated from the subsequent phase. The payoff structure of the subsequent phase that was used to quantify the update of risk-taking behavior was identical across the experimental conditions. The manipulated phase included baseline, positive, or negative experience, respectively. The length of initial experience was not manipulated, as it did not reliably modulate the change in risk taking in Experiment 1.

Although several hypotheses follow from previous literature, the analyses conducted in this study were exploratory. The following hypotheses were formulated after data collection. In Experiment 1, if initial experience were long, its profound and constraining effect on risk-taking behavior could be hypothesized (i.e., more/less balloon inflations after positive/negative experience, cf. ^8, 11^). In contrast, a more limited primacy effect could be expected if initial experience were short. In the latter case, initial and more recent experiences would be combined, which yields similar risk-taking behavior later in the task, irrespective of the valence of initial experience ^31^. In Experiment 2, it could be hypothesized that the effect of initial experience would be carried over to the subsequent task phase ^32, 33^. Since the separate manipulated phase of Experiment 2 consisted of the same number of balloons as the long condition of Experiment 1, similar results were expected to be found. Having been aware of the results of Experiment 1, after positive and negative experience in the initial phase, the presence of the primacy effect was hypothesized, which could be, at least partially, overcome (i.e., balloon inflations would be overall less/more after negative/positive experience but could increase/decrease as the task progresses). After the baseline experience, a relatively constant risk-taking behavior was expected that would be similar between the initial and subsequent task phases. In addition, it was also expected that negative experience would be more different from baseline than positive experience in terms of balloon inflations (cf. ^34^).

## Experiment 1

In Experiment 1, using a modified BART, we manipulated the first five or ten balloons out of the 30 balloons. We created lucky and unlucky runs of trials by providing the possibility to inflate the balloons to a larger size (lucky condition) or to experience frequent balloon bursts even after relatively few balloon pumps (unlucky condition). We crossed these conditions with the length of the initial experience (long condition: ten balloons vs. short condition: five balloons) to check whether the length of exposure to lucky or unlucky events differently influenced subsequent risk-taking behavior. These manipulations were based on the assumption that participants tended to gain some experience with the task in terms of action-outcome mapping (i.e., they tended to inflate the balloons instead of being risk averse, ^2, 35^), which would have enabled them to experience lucky and unlucky runs of events.

### Method

#### Participants

Eighty healthy young adults took part in Experiment 1, 20 in each experimental condition. As we used a between-subjects design, one participant performed only one experimental condition. This contrasts with previous studies testing the effect of initial experience or framing in the BART ^28, 29, 34, 36^. Although between-subjects designs involve known disadvantages (e.g., individual variability across experimental groups), they limit carryover effects that might originate from the complex experimental manipulation used here (i.e., luck and length). While the above-mentioned previous studies could not provide directly applicable effect size measures for the current between-subjects design, based on their findings, an f(U) effect size of at least 0.25 could be expected for the adjustment of the initial luck effect (i.e., interaction between task phase and condition). According to a power analysis with G*Power software version 3.1.9.4; ^37^, with an alpha level of 0.05 and a desired power level of 0.85, the required total sample size would be at least 60 participants. Therefore, our sample of 80 participants was sufficiently large to detect differences in the adjustment of risk-taking behavior across the conditions.

Participants were students from different universities in Budapest and young adult volunteers. All participants had normal or corrected-to-normal vision and none of them reported a history of any neurological and/or psychiatric condition. All of them provided written informed consent before enrollment. The experiment was approved by the United Ethical Review Committee for Research in Psychology (EPKEB) in Hungary and by the research ethics committee of Eötvös Loránd University, Budapest, Hungary; and it was conducted in accordance with the Declaration of Helsinki. As all participants were volunteers, they were not compensated for their participation. Descriptive characteristics of participants randomly assigned to the four different experimental conditions in Experiment 1 are presented in Table 1.

**Table 1.** Descriptive data of demographic variables and BART performance in Experiments 1–2 and in an explorative study presented in the Supplementary Material.

| Experiment | Explorative study | Experiment 1 |  |  |  | Experiment 2 |  |  |
| --- | --- | --- | --- | --- | --- | --- | --- | --- |
|  | – | Lucky long | Lucky short | Unlucky long | Unlucky short | Lucky | Unlucky | Baseline |
| Condition | – | *M*(*SD*) | *M*(*SD*) | *M*(*SD*) | *M*(*SD*) | *M*(*SD*) | *M*(*SD*) | *M*(*SD*) |
| *n* | 156 | 20 | 20 | 20 | 20 | 30 | 30 | 30 |
| Gender [Male/Female] | 26/130 | 11/9 | 9/11 | 11/9 | 5/15 | 16/14 | 14/16 | 18/12 |
| Age [years] | 21.4 (3.9) | 22.4 (1.6) | 21.0 (1.1) | 22.2 (1.3) | 21.8 (1.3) | 21.5 (1.6) | 21.6 (1.6) | 21.1 (1.4) |
| Education [years] | 14.6 (2.1) | 15.5 (1.3) | 14.2 (0.9) | 15.0 (1.1) | 15.2 (0.9) | 14.8 (1.4) | 14.9 (1.4) | 14.8 (1.6) |
| Mean adjusted pumps | 8.3 (2.3) | 8.7 (1.7) | 9.3 (1.3) | 5.7 (0.9) | 6.5 (0.9) | 8.1 (2.4) | 7.1 (1.4) | 7.9 (2.4) |
| Number of balloon bursts | 12.4 (4.1) | 11.3 (5.3) | 14.2 (4.3) | 10.1 (3.2) | 9.8 (2.2) | 12.1 (4.5) | 9.5 (3.2) | 11.0 (4.3) |
| Total score | 711.7 (256.4) | 797.2 (146.4) | 758.6 (127.1) | 405.1 (83.3) | 517.0 (91.3) | 651.1 (153.0) | 624.0 (153.8) | 644.6 (142.9) |

*Note.* In Experiment 1, BART variables were calculated for the whole task (30 balloons), which included the initial phase with experimental manipulation (biased balloon characteristics). In Experiment 2, BART variables were calculated for the subsequent phase (30 unbiased balloons).

#### Stimuli, design, and procedure

The appearance of the BART was the same as described in previous studies ^27, 38–41^. This version of the task was written in Presentation (v. 18.1, Neurobehavioral Systems). According to the instructions, participants were asked to achieve as high score as possible by inflating empty virtual balloons on the screen. They were also told that they were free to pump as much as they felt like, however, the balloon might burst. After each successful pump, the accumulated score on a given balloon (temporary bank) simultaneously increased with the size of the balloon. Instead of further pumping the balloon, participants could have finished the actual balloon trial and collected the accumulated score, which was transferred to a virtual permanent bank. Two response keys on a keyboard were selected either to pump the balloon or to finish the trial. There were two possible outcomes as results of a pump: The size of the balloon together with the score inside increased (positive feedback) or the balloon burst (negative feedback). The balloon burst ended the actual trial, and the accumulated score on that balloon were lost, but this negative event did not decrease the score in the permanent bank.

One point was added to the temporary bank for the first successful pump, two for the second (i.e., the accumulated score for a given balloon was 3), three for the third (i.e., the accumulated score was 6), and so on. Five information chunks persistently appeared on the screen during the task: (1) the accumulated score for a given balloon in the middle of the balloon, (2) the label “Total score” representing the score in the permanent bank, (3) the label “Last balloon” representing the score collected from the previous balloon, (4) the response key option for pumping the balloon and (5) the other response key option for collecting the accumulated score. After collecting the accumulated score and ending the balloon trial, a separate screen indicated the gained score. The feedback screen, indicating either the gained score or a balloon burst, was followed by the presentation of a new empty (small-sized) balloon denoting the beginning of the next trial.

Although the surface structure of the present task was consistent with the original version ^24^ and with those described in our previous studies ^27, 41^, the deep structure was modified. In the former versions, each successive pump not only increases the chance to gain more reward but also the probability of balloon burst and the accumulated reward to be lost. Therefore, the optimal choice is to inflate the balloon until a certain size, after that, further pumps are disadvantageous ^4^. This way, as Schonberg, et al. ^25^ pointed out, “increased risk is confounded with varying expected values” (p. 17) in the BART. In the present modified version, the probability of a balloon burst was zero until a certain threshold (maximum number of successful pumps) but it was one for the next pump (for a similar design, see ^42^). Although this design does not allow us to quantify how participants handle the dynamic change of burst probabilities within each balloon (across pumps), the risk level increases with each successive pump. Particularly, participants must consider the possibility to gain even more reward (the added points increase with each pump, see above) or to lose the already accumulated reward ^43, 44^.

Due to these modifications, the effect of positive and negative feedback and the sense of luckiness might become more emphasized, as balloon bursts could be experienced at various balloon sizes (see below). In all conditions, participants were not informed about the structure of the task and the change from the initial phase to the remaining phase in the maximum number of successful pumps. With repeated balloon inflations, they had the opportunity to experience the payoff structure. Therefore, as the original version, even this task measured decision making under uncertainty and decisions from experience ^4, 6^. We should nevertheless note that as the structure of the task considerably differed from that of the original BART, caution is needed when the current results are compared to those with the original task version.

Participants had to inflate 30 balloons in the task. We chose specific values for the *maximum number of successful pumps* on each of the 30 balloons. The maximum number of successful pumps (balloon tolerance) means that if a participant inflated the given balloon one pump further, the balloon burst. The maximum number of successful pumps for each balloon in each condition is presented in Table 2. These values were fixed across participants but differed across conditions.

**Table 2.**
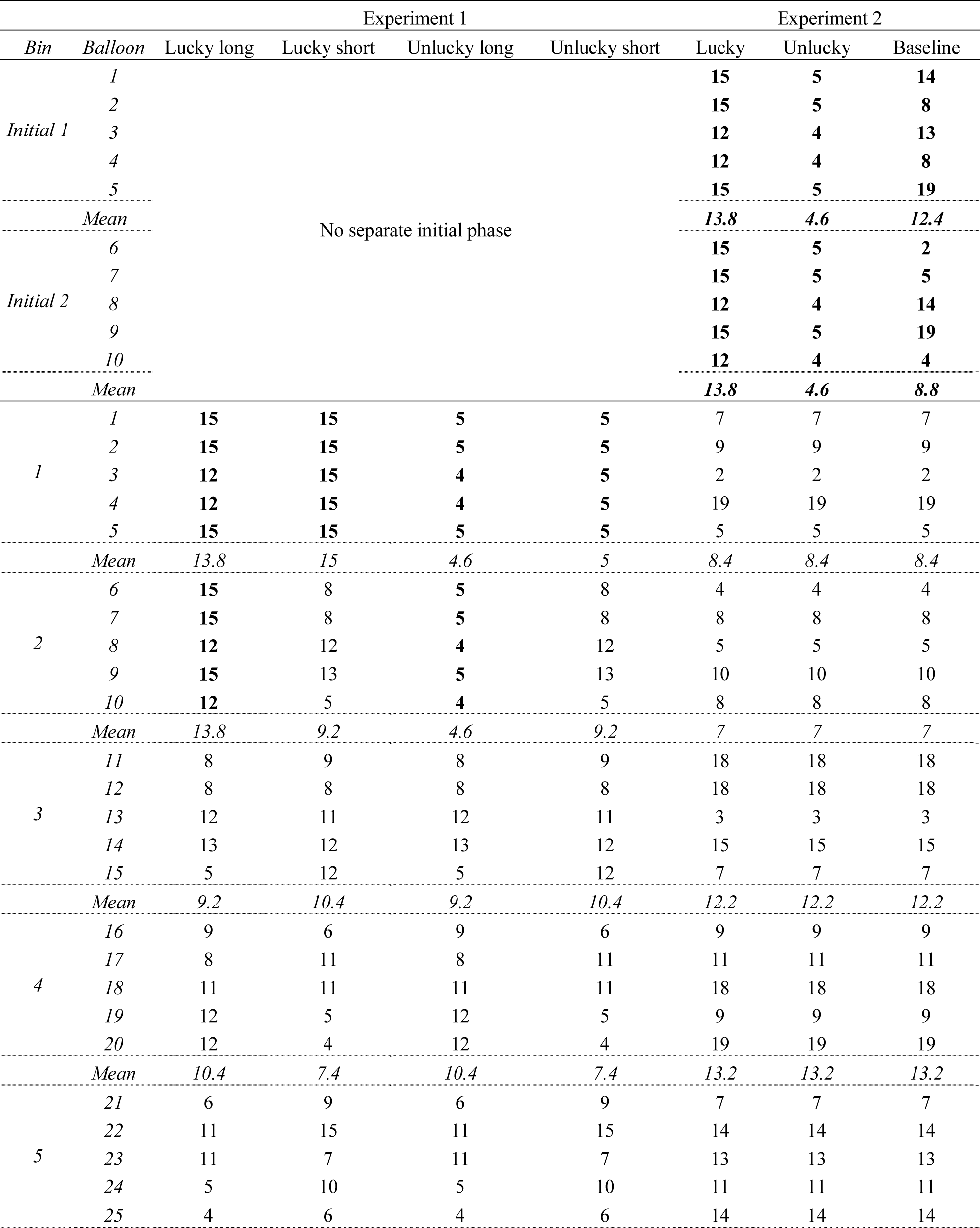
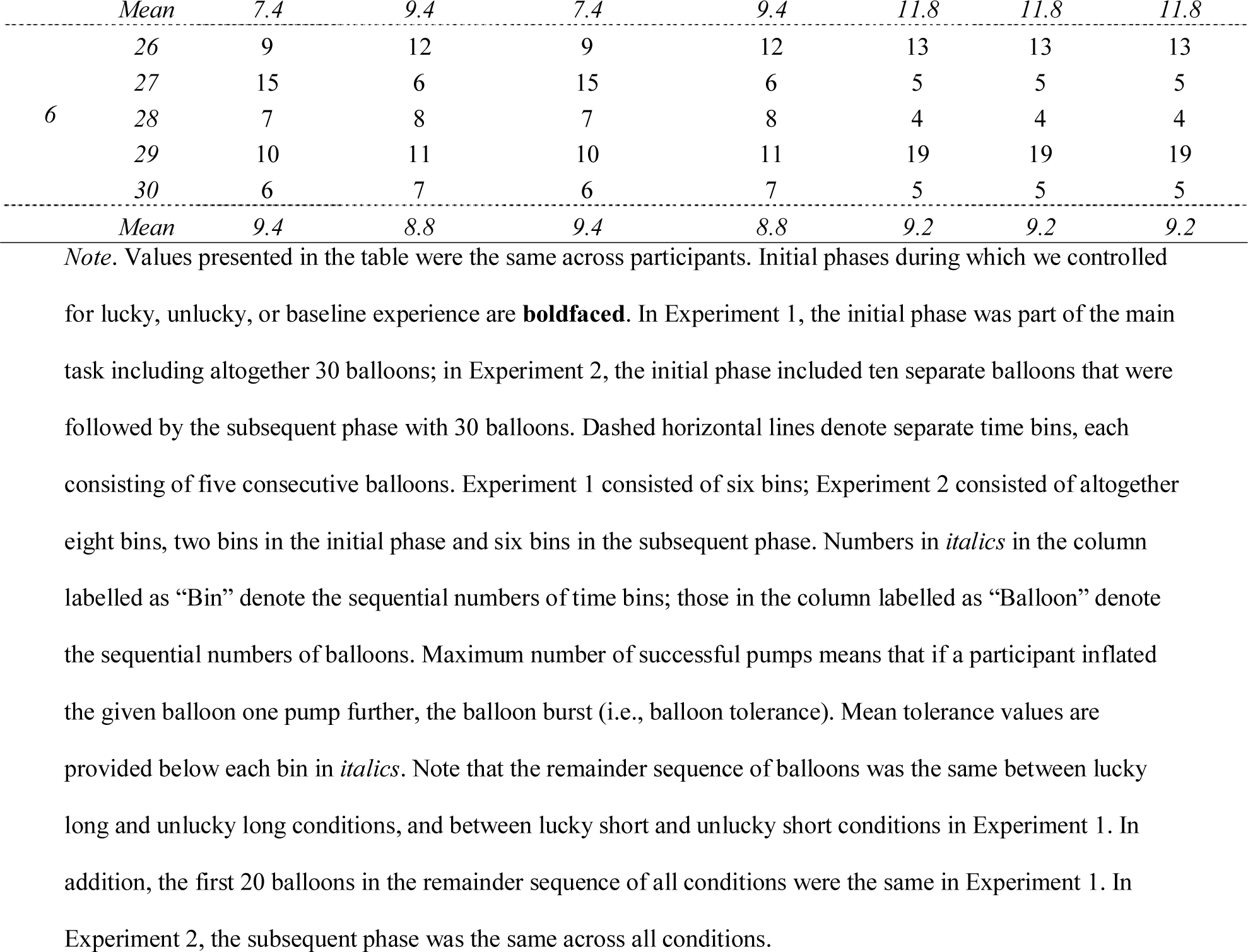
Maximum number of successful pumps (balloon tolerance) for each balloon in each condition in Experiments 1–2.

Regarding the initial balloons, in the *lucky long* condition, the maximum number of successful pumps for each of the first ten balloons was determined by a fixed variation of 12 and 15. In the *lucky short* condition, the maximum number of successful pumps for each of the first five balloons was 15 (same for all five initial balloons). In the *unlucky long* condition, for each of the first ten balloons, this was a fixed variation of four and five. In the *unlucky short* condition, for each of the first five balloons, this was five (same for all five initial balloons).

We also determined the maximum number of successful pumps for each balloon in the remaining 20 or 25 balloons (after initial balloons): These were identical to one another between the equal-length conditions (i.e., lucky vs. unlucky long conditions and lucky vs. unlucky short conditions, see Table 2) and across participants, as indicated above. The fixed series of values were a random sequence of integers generated from a uniform distribution between three and 15 in Matlab 8.5 (MathWorks Inc.). We had chosen this interval between minimum and maximum values to ensure that the remainder of the experiment was different from the initial phase and balloons could be, on average, inflated up to a “medium” size.

There was no significant difference in the mean of the maximum number of successful pumps – calculated for the whole task – between the lucky long (*M* = 10.66, *SD* = 3.44) and lucky short (*M* = 10.03, *SD* = 3.47) conditions (*p* = .412) and between the unlucky long (*M* = 7.60, *SD* = 3.28) and unlucky short (*M* = 8.36, *SD* = 3.05) conditions (*p* = .309). Hence, the length of the initial phase did not change the overall potential of risk taking. Lucky long and unlucky long conditions differed significantly (*p* < .001) in the mean of the maximum number of successful pumps, so did lucky short and unlucky short conditions (*p* = .023).

To test whether participants used any task-solving strategies and gained awareness about the regularities underlying the BART, a short verbal interview with two questions was administered by the experimenters immediately after finishing the task. We asked participants (1) how they solved the task, how they tried to achieve as high score as possible; and (2) whether they have noticed any regularity in the sequence of balloon bursts. The experimenters were not blind to the research question or condition of each participant. The sixth author (E. T-F. who was not blind to the research question or condition either) rated participants’ answers for the following aspects: (1) mentioning change between task phases, (2) mentioning any systematic task-solving approach throughout the task, irrespective of its adaptiveness, (3) mentioning beliefs reflecting the so-called gambler’s fallacy, (4) and mentioning beliefs in the presence of lucky/unlucky series. Independent raters were not involved. Altogether, the interviews were not conducted and evaluated according to a standardized protocol. Therefore, the related results cannot provide accurate insights into task-solving strategies, because of which these results are not reported here.

#### Data analyses

To evaluate the effect of initial experience on risk-taking behavior, we performed linear mixed-effects analyses on the number of pumps on each balloon that did not burst as the dependent variable. This adjusted pump number is a conventionally used index in the BART literature ^24, 28, 45, 46^. Although accumulating evidence indicates that risk-taking propensity or risk preferences measured by self-report questionnaires and by behavioral measures are poorly correlated ^12, 42, 47^, in line with the BART literature, we considered the adjusted pump number as a measure of risk taking (cf. ^48^). The advantage of using linear mixed-effects models is that they could account for the nonindependence of observations nested within participants (i.e., balloon pumps are repeated measures observations within participants). In addition, the dependent variable (number of pumps) does not have to be aggregated at the level of participants, which could yield less unexplained variance and higher statistical power.

To determine more clearly how long the effect of initial experience would persist, each balloon was assigned to a five-balloon-long time bin of the task. Although dividing the task into five-balloon-long bins might seem arbitrary, it is in accordance with the length of each manipulation (equal to the short ones, half of the long ones). We found easier to interpret the results based on bin-wise analyses that has also been frequent in the BART literature (e.g., ^2, 29, 49^). Therefore, further linear mixed-effects analyses were performed on the number of pumps on balloons that did not burst while considering the balloons’ bin-wise assignment. Importantly, to account for the effect of time bin, no aggregation of the dependent variable (number of pumps) was needed at the level of bins.

The balloon-wise and bin-wise linear mixed-effects analyses were performed on the number of pumps on *remaining* (unbiased) balloons that did not burst in the equal-length conditions. Thus, separate models were fit to the data of the long-manipulation conditions (lucky and unlucky long) and that of the short-manipulation conditions (lucky and unlucky short). In the long-manipulation conditions, the remaining phase of the task consisted of 20 balloons or four bins, in the short-manipulation conditions, the remaining phase consisted of 25 balloons or five bins. The tolerances of the remaining balloons were the same across the equal-length conditions, but these values were shifted across the different-length conditions. (Particularly, tolerance values were the same between Balloons 11–30 in the long conditions and Balloons 6–25 in the short conditions, see Table 2). Therefore, the possibility to inflate a balloon during the remaining phase was differently constrained across the conditions, which warranted the separate analyses of the temporal change in risk-taking behavior after the end of initial manipulation. In addition to these analyses, a linear mixed-effects model including all experimental factors was also fit to the *entire task* with 30 (manipulated and unbiased) balloons. (Note that the bin-wise version of this model is presented only in Supplementary Table S1.) The latter analysis aimed to test the potential effect of length manipulation on pumping behavior and its adjustment.

All analyses were performed using the *lmer* function implemented in the *lme4* package ^50^ of R ^51^. The *p*-values for fixed effects were computed using Satterthwaite’s degrees of freedom method with the *lmerTest* package ^52^. The schematic structures of the six linear models are summarized below with the dependent variable on the left side of the “∼” symbol and the predictor variables on the right side:

Model 1: Number of pumps on *remaining* balloons that did not burst in the long-manipulation conditions ∼ Luck, Balloon, Luck * Balloon + (1 | participant)

Model 2: Number of pumps on *remaining* balloons that did not burst in the long-manipulation conditions ∼ Luck, Bin, Luck * Bin + (1 | participant) (*Bin with 4 levels*)

Model 3: Number of pumps on *remaining* balloons that did not burst in the short-manipulation conditions ∼ Luck, Balloon, Luck * Balloon + (1 | participant)

Model 4: Number of pumps on *remaining* balloons that did not burst in the short-manipulation conditions ∼ Luck, Bin, Luck * Bin + (1 | participant) (*Bin with 5 levels*)

Model 5: Number of pumps on *all* balloons that did not burst ∼ Luck, Length, Balloon, Luck * Length, Luck * Balloon, Length * Balloon, Luck * Length * Balloon + (1 | participant)

The factors *Luck* (lucky vs. unlucky), *Length* (long vs. short), *Balloon* or *Bin*, and their *two-way* and *three-way* interactions were entered as fixed effects into the models. The *Luck*, *Length*, and *Bin* factors were treated as categorical predictors and *Balloon* as a continuous predictor. Participants were modeled as random effects (random intercepts). As we used the treatment contrast, the reference level of a given factor served as the baseline to estimate the other levels. These were *lucky*, *long*, *Bin2* (Model 4), and *Bin3* (Model 2). The models were fit with restricted maximum likelihood parameter estimates (*REML*). Pairwise comparisons were performed by the *glht* function ^53^ (*p* values were adjusted with single-step method).

### Results

To ease meta-analytic work, Table 1 shows overall performance on the different versions of the BART presented in this paper measured by classical indices such as the mean adjusted number of pumps, the number of balloon bursts, and the total score that is the sum of reward (points) in the end of the task. In the results section below, the “number of pumps” or “pump number” refers to the number of pumps on balloons that did not burst. Only those effects are detailed from the balloon-wise and bin-wise analyses that are relevant regarding the hypotheses. The summary of all effects included in the linear mixed-effects models is presented in Table 3. Since the treatment contrast provides inferential tests of simple effects and simple interactions (except for the highest-order effects), these terms are used below, if relevant.

**Table 3.**
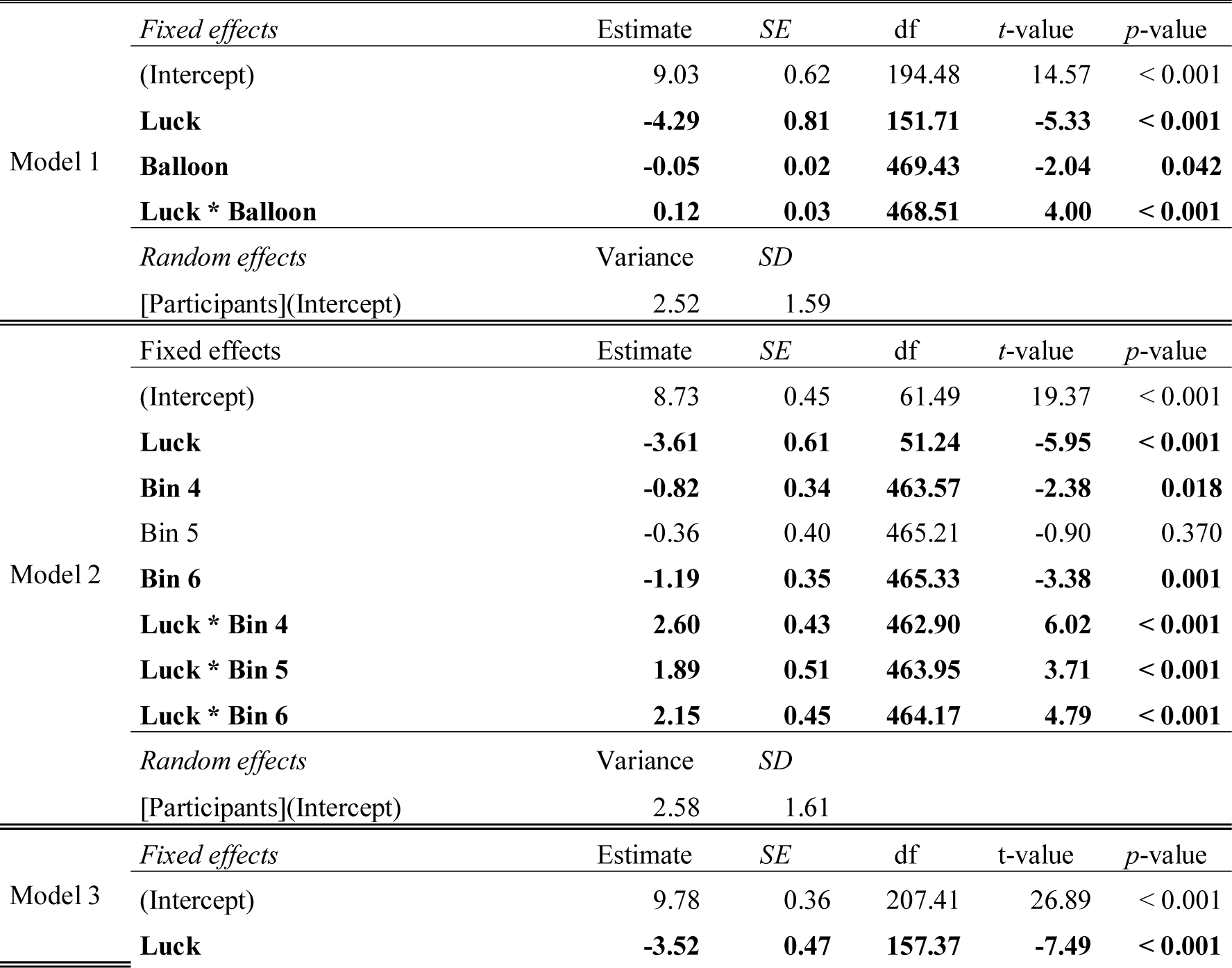
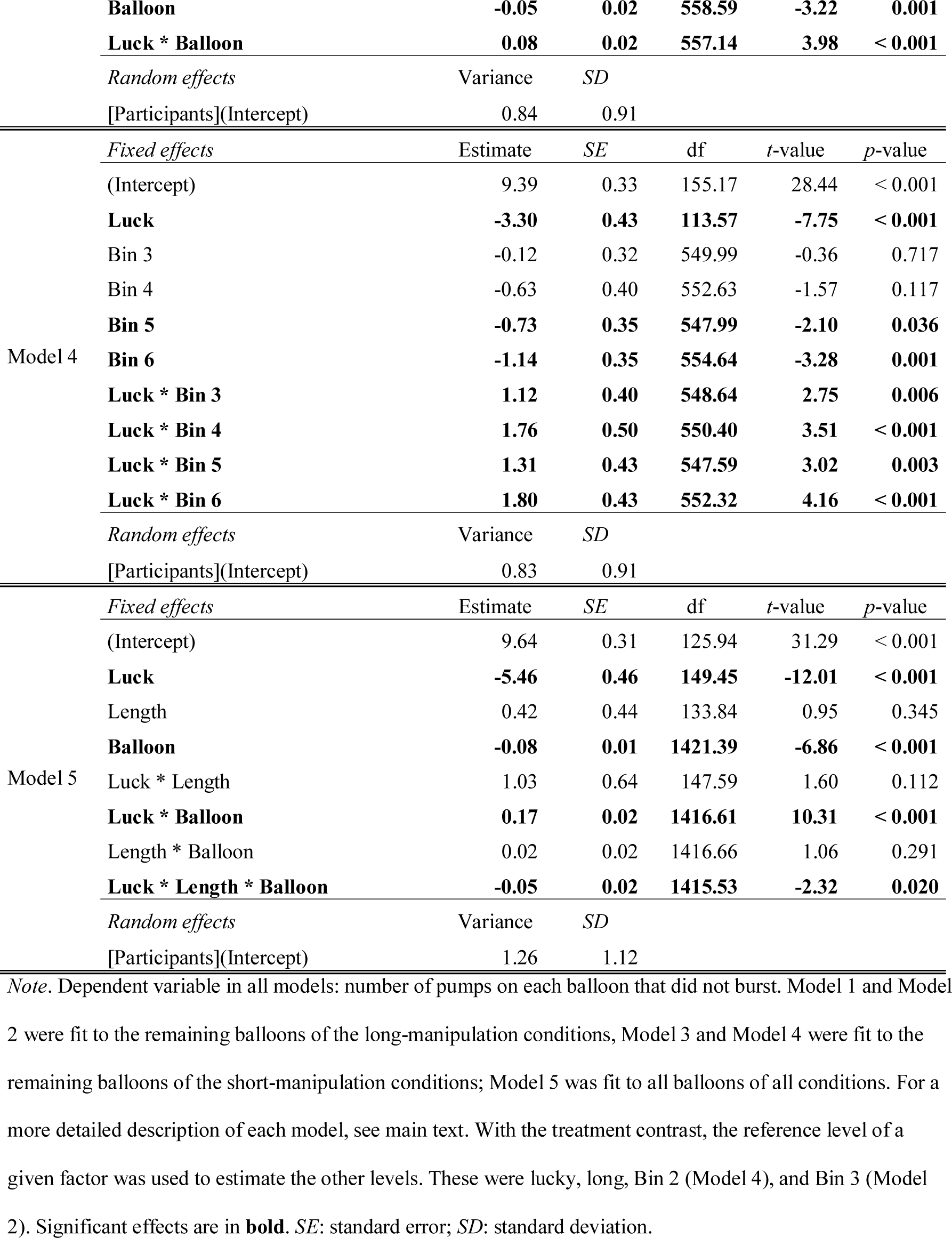
Summary of linear mixed-effects models on the number of pumps that did not burst in Experiment 1.

#### Risk-taking behavior in the lucky long and unlucky long conditions (Model 1 & Model 2)

After *long* initial manipulation had ended, unlucky experience yielded lower number of pumps on the remaining balloons than lucky experience (significant simple effect of *Luck*, β = −4.29, *SE* = 0.81, *t*(151.71) = −5.33, *p* < .001). However, as the task progressed, while the number of pumps increased on the remaining balloons in the unlucky long condition, it decreased in the lucky long condition (significant *Luck * Balloon* interaction, β = 0.12, *SE* = 0.03, *t*(468.51) = 4.00, *p* < .001; see Fig. 1a, c). More specifically, as compared with the first Bin after initial manipulation (i.e., Bin3), a significant decrease in the lucky long vs. unlucky long difference (*Luck* effect) was evidenced from Bin4 until the end of the task (all βs ≥ 1.89, *SE*s ≤ 0.51, *t*s ≥ 3.71, *p*s < .001, see Table 3). A contrast matrix was set up that defined the comparison of *Luck* effect across the levels of *Bin*. Pairwise tests showed that the *Luck* effect did not change further across Bin4–Bin6 (│*z*s│ ≤ 1.46, *p*s ≥ .310).

**Figure 1.**
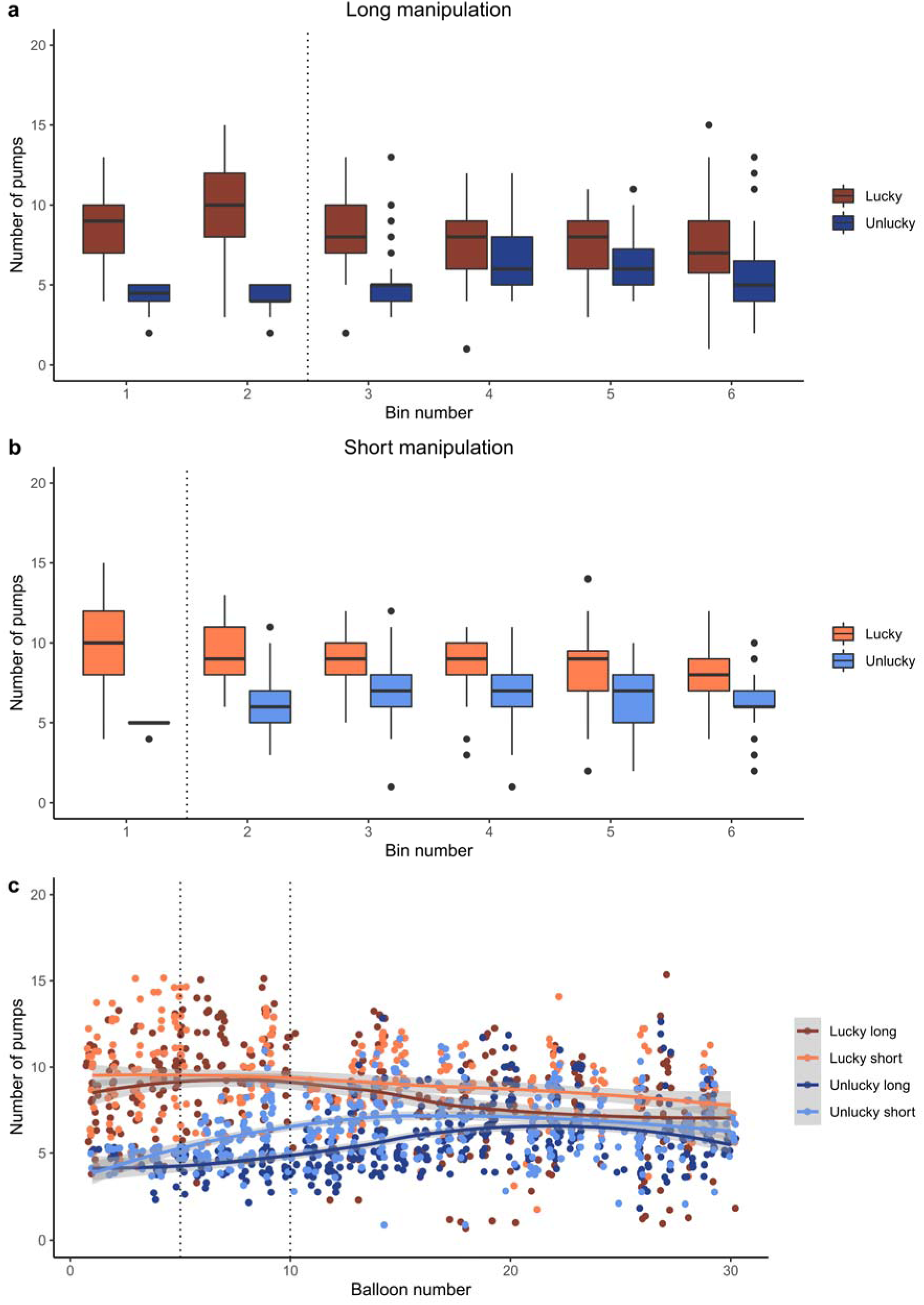
Results of Experiment 1. The numbers of pumps on balloons that did not burst are presented in each time bin in the long manipulation (a) and short manipulation (b) conditions, respectively. Vertical dotted lines indicate that (lucky or unlucky) initial manipulation ended after the second (long manipulation) or the first (short manipulation) time bin. Each time bin consists of five consecutive balloons. Figure part (c) shows the numbers of pumps on balloons that did not burst across all conditions and balloons. Shaded areas represent 95% confidence intervals for means, depicted with continuous lines. Points denote individual data points in all figure parts.

#### Risk-taking behavior in the lucky short and unlucky short conditions (Model 3 & Model 4)

After *short* initial manipulation had ended, unlucky experience yielded lower number of pumps on the remaining balloons than lucky experience (significant simple effect of *Luck*, β = −3.52, *SE* = 0.47, *t*(157.37) = −7.49, *p* < .001). Meanwhile, the number of pumps increased on the remaining balloons in the unlucky short condition and decreased in the lucky short condition (significant *Luck * Balloon* interaction, β = 0.08, *SE* = 0.02, *t*(557.14) = 3.98, *p* < .001; see Fig. 1b, c). Particularly, as compared with the first Bin after initial manipulation (i.e., Bin2), a significant decrease in the lucky short vs. unlucky short difference (*Luck* effect) appeared in Bin3 and was present until the end of the task (all βs ≥ 1.12, *SE*s ≤ 0.50, *t*s ≥ 2.75, *p*s ≤ .006, see Table 3). Pairwise tests showed that this *Luck* effect did not change further across Bin3–Bin6 (│*z*s│ ≤ 1.73, *p*s ≥ .304).

#### Interim summary

Due to both long and short initial manipulations, the number of pumps was consistently lower in unlucky than in lucky conditions across the remaining balloons of the task. Meanwhile, this difference decreased in the second half of the task after experiencing approximately five unbiased balloons following initial manipulation. While participants in the unlucky condition increased pumping, those in the lucky condition slightly decreased this behavior in the remainder of the task, indicating the adjustment of behavior to recent experience. Although it seems that participants in all conditions adjusted their behavior in an opposite manner as a function of luck, it was not clear from these analyses whether the effect of long or short initial experience was more pronounced. This was tested below with Model 5.

#### Risk-taking behavior across all conditions and balloons (Model 5)

When all conditions and balloons (manipulated and unbiased) were considered together, in line with the results above, consistently lower pump number in the unlucky than in the lucky condition was found (significant simple effect of *Luck*, β = −5.46, *SE* = 0.45, *t*(149.45) = −12.01, *p* < .001). The simple effect of *Length* was not significant, β = 0.42, *SE* = 0.44, *t*(133.84) = 0.95, *p* = .345 (but see the bin-wise model in Supplementary Table S1). As shown by the significant *Luck * Length * Balloon* interaction, β = −0.05, *SE* = 0.02, *t*(1415.53) = −2.32, *p* = .020, although the number of pumps increased in the unlucky condition and decreased in the lucky condition (significant simple interaction of *Luck * Balloon*), this change differed between the long and short conditions (see Fig. 1c).

Based on the bin-wise analysis summarized in Supplementary Table S1 and the visual inspection of Figure 1c, this significant three-way interaction indicated the local effectiveness of initial manipulation and not the length of initial experience modulating behavior adjustment. Particularly, the lucky vs. unlucky difference was smaller in the short than in the long manipulation conditions in Bin2 and Bin3 than at the beginning of the task. This was due to short manipulation had ended by Bin2 and the lucky vs. unlucky difference started to decrease (see the steeply increasing pump number in the unlucky short condition). More importantly, during the second half of the task (Bin4, Bin5, and Bin6 as compared with Bin1), no difference appeared between the short and long conditions in how the lucky vs. unlucky difference changed as the task progressed. Therefore, experiencing unlucky balloons either for longer or shorter duration yielded similar pump number after the integration of recent experience. Altogether, these modeling results with all experimental factors considered not only corroborated previous results (Model 1–4) but also revealed that behavior adjustment in the second half of the task was comparable between the length manipulation conditions.

### Discussion

In Experiment 1, we showed that initial experience influenced subsequent risk-taking behavior. Unlucky compared with lucky initial experience led to overall lower risk taking. More importantly, approximately five balloons after the change in structure, participants with both lucky and unlucky experience started to update their behavior: The number of pumps increased following unlucky experience, while it decreased following lucky experience in the remainder of the task. The pattern of change was comparable between long and short manipulation conditions. This is partially in contrast to the expectation that a persistent risk-averse behavior would be present after a long series of unlucky events (i.e., primacy effect). Namely, albeit unlucky participants converged to a more risk-averse behavior in both length manipulation conditions, behavior adjustment was still observed due to combining the outcomes of more recent balloon pumps with initial experience.

In Experiment 1, the number of balloons and their tolerance in the remaining phase of the task differed across conditions, which could have influenced how recent outcomes were evaluated and used in updating. Furthermore, one cannot tell whether lucky and unlucky experiences were more positive or more negative than a baseline experience. Experiment 2 aimed to address these shortcomings. Since the length of initial experience did not modulate behavior adjustment, length was not manipulated in Experiment 2. Without length manipulation, it was also ensured that all participants encounter the same number of balloons. Therefore, any change in risk-taking behavior would be due to the valence of initial experience and not merely a longer horizon for post-manipulation updating.

## Experiment 2

In Experiment 2, we manipulated a separate initial phase with ten balloons that preceded a subsequent phase with 30 balloons. This design enabled us to analyze risk-taking behavior on the exact same sequence of 30 balloons across all conditions in the subsequent phase. In the initial phase, we used three conditions: baseline, lucky, and unlucky. With the use of the baseline condition, we could compare with a reference level whether individuals are more prone to take risks due to lucky initial experience and less prone to take risks due to unlucky initial experience.

### Method

#### Participants

Ninety young adults took part in Experiment 2, 30 in each experimental condition (between-subjects design). None of them participated in Experiment 1. According to a similar power analysis as in relation to Experiment 1, the required total sample size would be at least 63 participants. Thus, our sample of 90 participants was sufficiently large to detect differences in the adjustment of risk-taking behavior across the conditions.

Participants were students from different universities in Budapest and young adult volunteers. All participants had normal or corrected-to-normal vision and none of them reported a history of any neurological and/or psychiatric condition. All of them provided written informed consent before enrollment. The experiment was approved by the United Ethical Review Committee for Research in Psychology (EPKEB) in Hungary and by the research ethics committee of Eötvös Loránd University, Budapest, Hungary; and it was conducted in accordance with the Declaration of Helsinki. As all participants were volunteers, they were not compensated for their participation. Descriptive characteristics of participants randomly assigned to the three different conditions are presented in Table 1.

#### Stimuli, design, and procedure

The appearance of the task was the same as in Experiment 1. Participants had to inflate altogether 40 balloons in this task. The first ten balloons belonged to the *initial phase*; the remaining 30 balloons belonged to the *subsequent phase*. Similar to Experiment 1, we chose specific values for the maximum number of successful pumps on each of the 40 balloons. The maximum pump values for each balloon in both phases, separately for each condition, are presented in Table 2.

Regarding the initial phase, in the *lucky* condition, we used the same fixed series of values as in the lucky long condition of Experiment 1 (either 12 or 15 pumps). Similarly, in the *unlucky* condition, we used the same fixed series of values as in the unlucky long condition of Experiment 1 (either four or five pumps). For the *baseline* condition, we generated a random sequence of ten integers from a uniform distribution between two and 19, as balloon burst was enabled only after the third pump, and the maximum number of successful pumps was 19 in those versions that were used in our previous studies (e.g., ^27^). Again, this series of baseline pump values was fixed and the same across participants. The mean of the maximum number of successful pumps – calculated for the initial phase – significantly differed across the three conditions, *F*(2, 18) = 17.55, *p* = .002. It was significantly lower in the unlucky than in the baseline and lucky conditions (unlucky: *M* = 4.60, *SD* = 0.52, baseline: *M* = 10.60, *SD* = 6.08, lucky: *M* = 13.80, *SD* = 1.55; unlucky vs. baseline: *p* = .012, unlucky vs. lucky: *p* < .001); but it was not significantly higher in the lucky than in the baseline condition (*p* = .131).

Regarding the 30 balloons of the subsequent phase, the fixed series of values were identical across the different conditions and across participants (see Table 2). These values were, again, generated from a uniform distribution between two and 19. Since the subsequent phase was the same but the initial phase was different, the mean of the maximum number of successful pumps – calculated for the initial and the subsequent phases together (whole task) – significantly differed across the three conditions, *F*(2, 78) = 7.72, *p* = .001 (unlucky: *M* = 8.88, *SD* = 5.26, baseline: *M* = 10.38, *SD* = 5.47, lucky: *M* = 11.18, *SD* = 4.92). However, the mean of the maximum number of successful pumps over the course of the whole task was not significantly higher in the lucky than in the baseline condition (lucky vs. baseline: *p* = .127; unlucky vs. baseline: *p* = .020; unlucky vs. lucky: *p* = .001).

Participants were told that they were going to have ten balloon trials in the initial phase of the task and then 30 balloon trials in the subsequent phase. The starting score was zero in both phases, about which participants were also informed. Importantly, no information was provided about the change in the payoff structure. The short interview as described in Experiment 1 was also administered after the task but was not analyzed in depth.

#### Data analyses

We followed the same approach as in Experiment 1. We performed linear mixed-effects analyses on the number of pumps on each balloon that did not burst as the dependent variable. Again, each balloon was assigned to a five-balloon-long time bin of the task. First, to check the effectiveness of initial manipulation, two models (balloon-wise and bin-wise) were fit to the data of the initial phase. Second, to evaluate the effect of initial experience on subsequent risk-taking behavior, two models were fit to the data of the subsequent phase.

The initial phase consisted of ten balloons or two bins; the subsequent phase consisted of 30 balloons or six bins. The factors *Condition* (baseline, lucky, unlucky), *Balloon* or *Bin*, and their *two-way interactions* were entered as fixed effects into the models. The *Condition* and *Bin* factors were treated as categorical predictors and *Balloon* as a continuous predictor. Participants were modeled as random effects (random intercepts). With the treatment contrast, the reference level of a given factor (i.e., *baseline, Bin1*) was used to estimate the other levels. The four linear models are summarized below with the dependent variable on the left side of the “∼” symbol and the predictor variables on the right side:

Model 1: Number of pumps on balloons that did not burst in the *initial phase* ∼ Condition, Balloon, Condition * Balloon + (1 | participant)

Model 2: Number of pumps on balloons that did not burst in the *initial phase* ∼ Condition, Bin, Condition * Bin + (1 | participant) (*Bin with 2 levels*)

Model 3: Number of pumps on balloons that did not burst in the *subsequent phase* ∼ Condition, Balloon, Condition * Balloon + (1 | participant)

Model 4: Number of pumps on balloons that did not burst in the *subsequent phase* ∼ Condition, Bin, Condition * Bin + (1 | participant) (*Bin with 6 levels*)

### Results

The “number of pumps” or “pump number” below refers to the number of pumps on balloons that did not burst. Only those effects are detailed from the balloon-wise and bin-wise analyses that are relevant regarding the hypotheses. The summary of all effects included in the linear mixed-effects models is presented in Table 4.

**Table 4.**
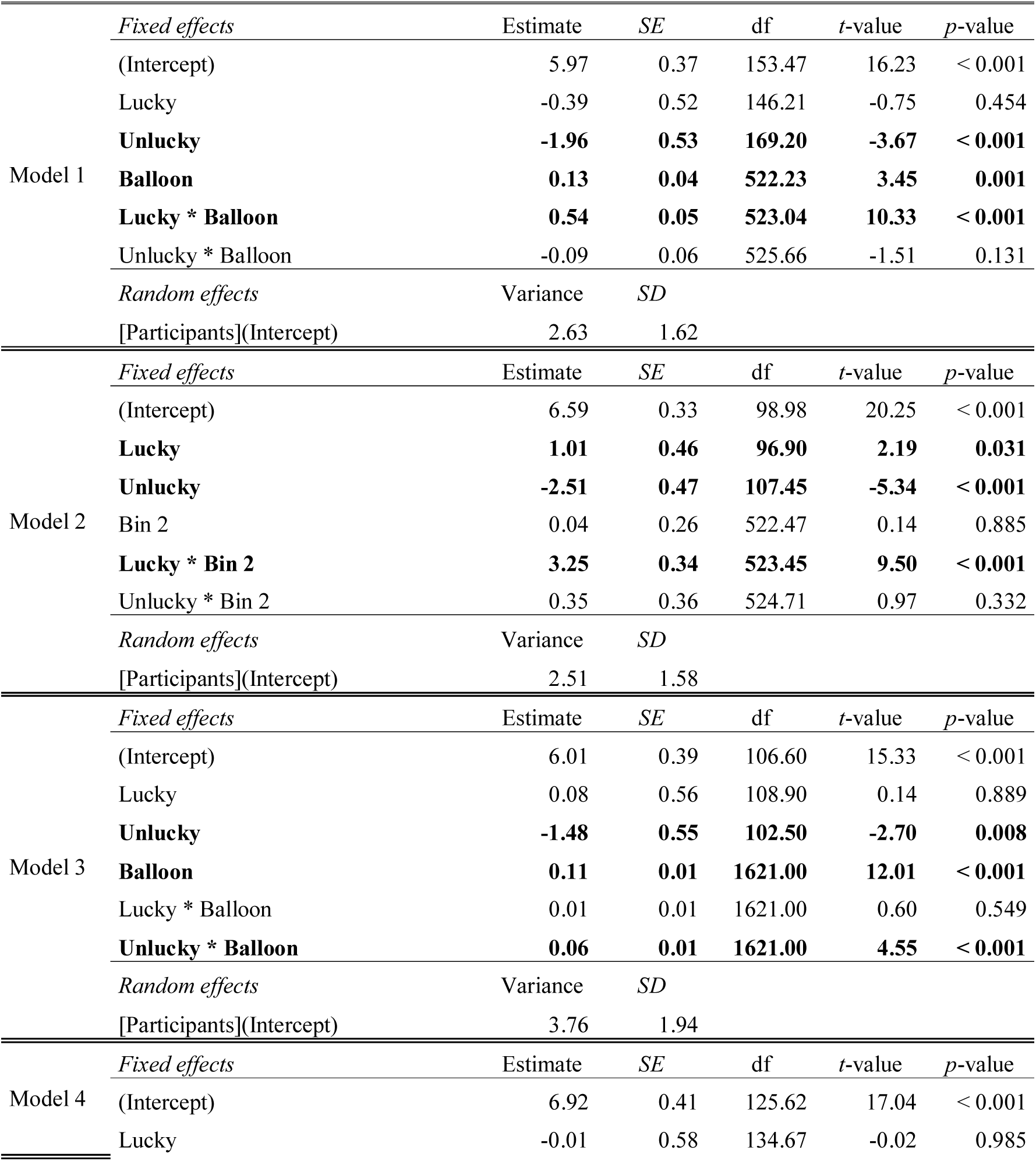
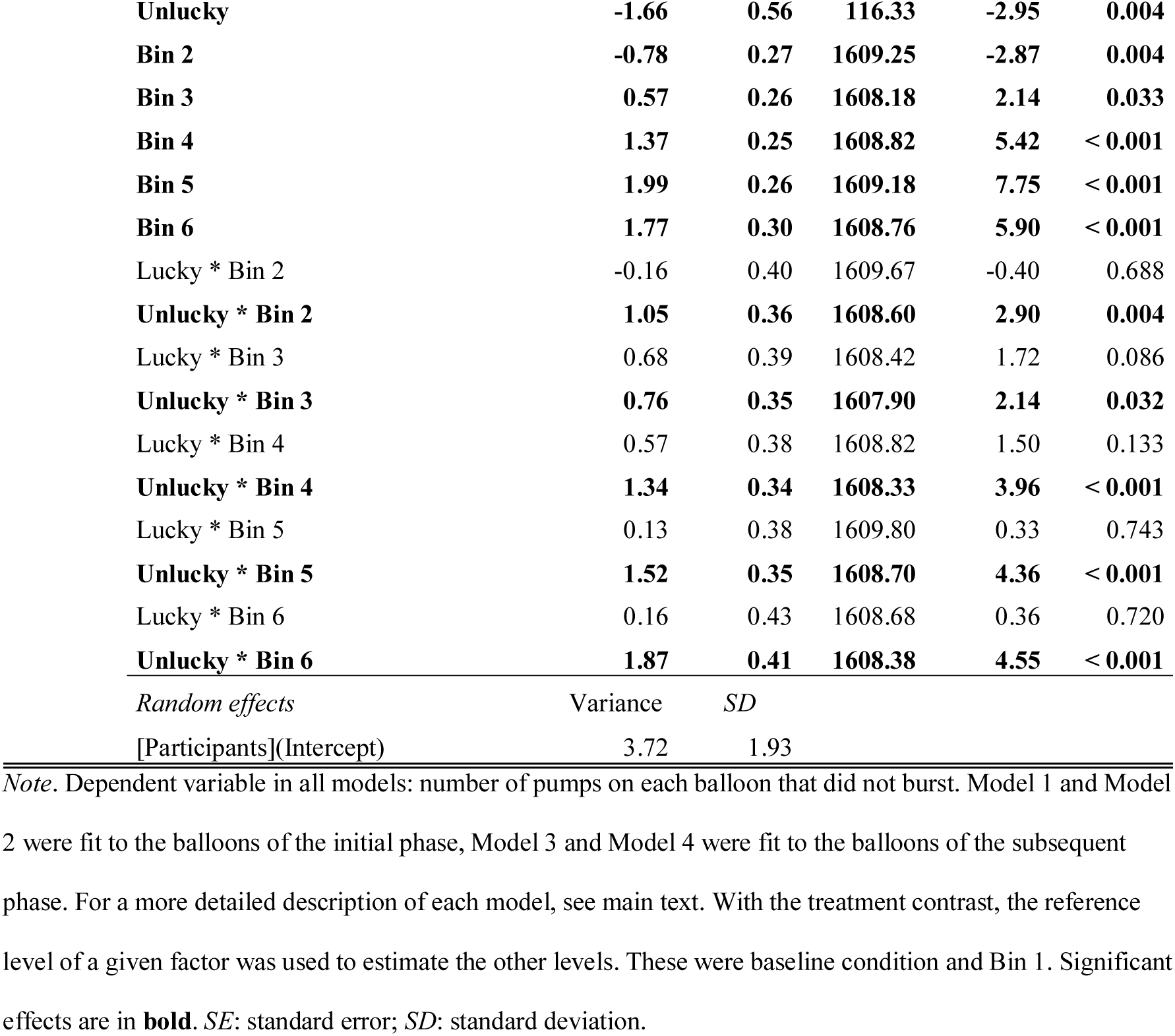
Summary of linear mixed-effects models on the number of pumps that did not burst in Experiment 2.

#### Risk-taking behavior in the initial task phase (Model 1 & Model 2)

Unlucky initial experience yielded significantly lower pump number on balloons of the *initial phase* than baseline experience (β = −1.96, *SE* = 0.53, *t*(169.20) = −3.67, *p* < .001). Lucky initial experience yielded significantly higher pump number than baseline experience only in the bin-wise model, β = 1.01, *SE* = 0.46, *t*(96.90) = 2.19, *p* = .031, and not in the balloon-wise one. However, participants in the lucky condition showed increased pump number relative to the baseline condition with later balloons, especially in Bin2 (significant *Condition * Balloon* and *Condition * Bin* interactions, β = 3.25, *SE* = 0.34, *t*(523.45) = 9.50, *p* < .001, see also Table 4 and Fig. 2a). Overall, initial manipulation seemed to be effective, as lucky and unlucky conditions differed from the baseline in the expected directions.

**Figure 2.**
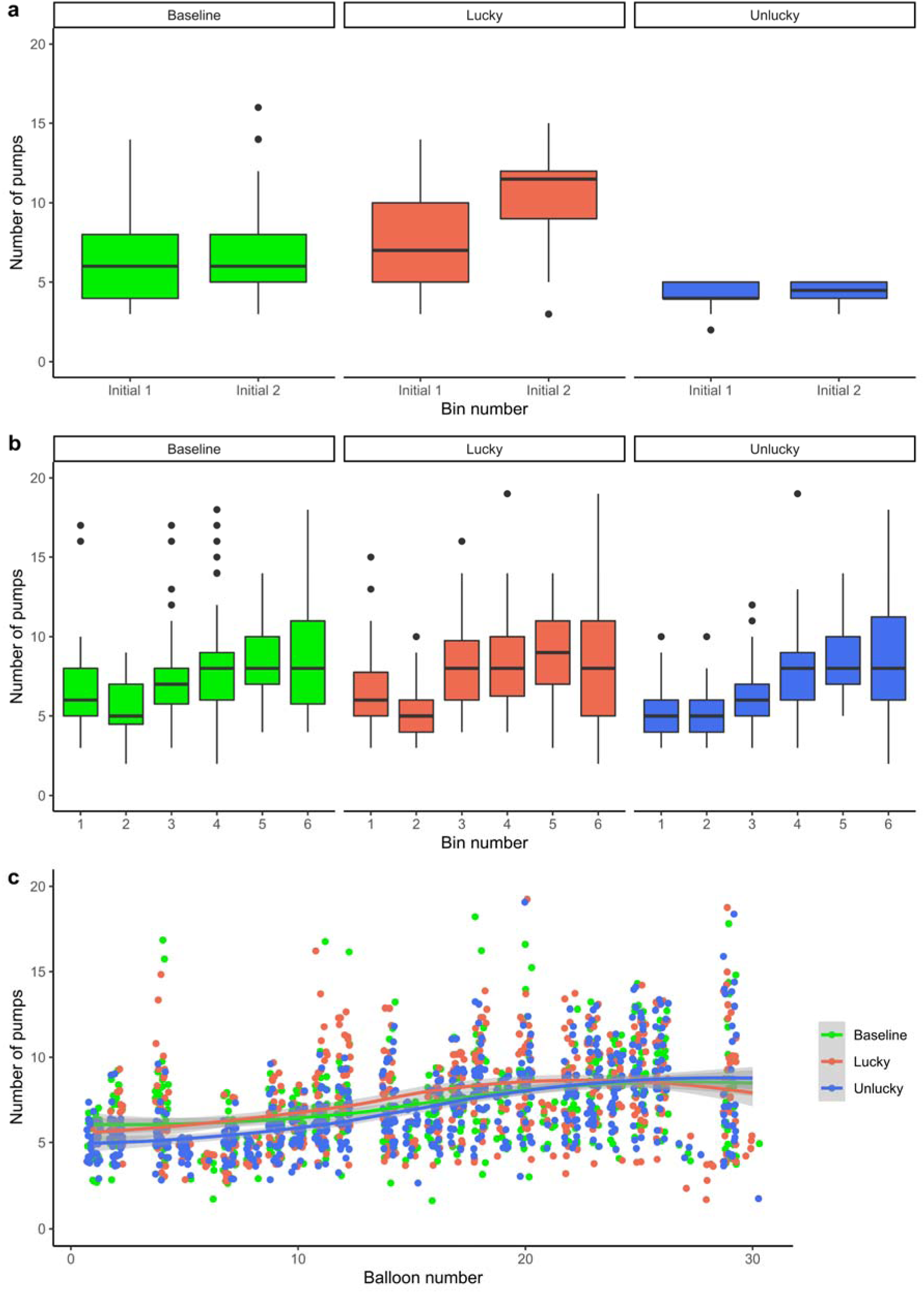
Results of Experiment 2. The numbers of pumps on balloons that did not burst are presented separately for each condition (baseline, lucky, unlucky) in each time bin of the initial (a) and subsequent (b) phases of the task. These phases were separated by a short pause lasting for a few seconds. Each time bin consists of five consecutive balloons. Figure part (c) shows the numbers of pumps on balloons that did not burst across all conditions and balloons. Shaded areas represent 95% confidence intervals for means, depicted with continuous lines. Points denote individual data points in all figure parts.

#### Risk-taking behavior in the subsequent task phase (Model 3 & Model 4)

Unlucky initial experience yielded significantly lower pump number on balloons of the *subsequent phase* than baseline experience (β = −1.48, *SE* = 0.55, *t*(102.5) = −2.70, *p* = .008). Meanwhile, the effect of lucky experience did not reliably differ from that of the baseline experience during the subsequent phase (β = 0.08, *SE* = 0.56, *t*(108.9) = 0.14, *p* = .889). Participants in all conditions increased pumping as the task progressed (simple effect of *Balloon*, β = 0.11, *SE* = 0.01, *t*(1621) = 12.01, *p* < .001; see Fig. 2b, c). As compared with Bin1, this increase was significant from Bin3 until the end of the task (all βs ≥ 0.57, *SE*s ≤ 0.30, *t*s ≥ 2.14, *p*s ≤ .033). Though, further change was not observed in the second half of the task, except between Bin4 and Bin5 (*z* = 2.81, *p* = .044; Bin4 vs. Bin6: *z* = 1.49, *p* = .628, Bin5 vs. Bin6: *z* = −0.80, *p* = .959).

The increasing pump number with further balloons differed between the unlucky and baseline conditions, β = 0.06, *SE* = 0.01, *t*(1621) = 4.55, *p* < .001, but it did not between lucky and baseline conditions, β = 0.01, *SE* = 0.01, *t*(1621) = 0.60, *p* = .549. This *Condition * Balloon* interaction was corroborated by the bin-wise analysis, indicating significantly increased pump numbers in the unlucky condition in all time bins relative to Bin1 (all βs ≥ 0.76, *SE*s ≤ 0.41, *t*s ≥ 2.14, *p*s ≤ .032; see Fig. 2b). At the same time, pairwise tests showed that across the levels of *Bin*, this relative increase in the unlucky condition did not change further (│*z*s│ ≤ 2.79, *p*s ≥ .063). Relative to Bin1, the unlucky vs. baseline difference was larger than the lucky vs. baseline difference in Bin2, Bin5, and Bin6, indicating a steeper increase in the unlucky than in the baseline or lucky conditions (*z*s ≥ 3.20, *p*s ≤ .017).

Overall, after completing approximately ten balloons of the subsequent phase, participants started to increase their pumping behavior that became consistent in the second half of the task. While this increase was similar in the baseline and lucky conditions, it was steeper in the unlucky condition, as those participants showed lower pump number in the first half of the task because of initial manipulation.

### Discussion

Experiment 2 involved not only lucky and unlucky conditions but also a baseline condition; and the initial phase with experimental manipulation was separated from the subsequent phase where the adjustment of risk-taking behavior was measured. Based on previous literature and the findings of Experiment 1, a primacy effect with a later behavioral shift was expected, meaning that lucky initial experience would have been related to decreased risk taking and unlucky initial experience would have been related to increased risk taking on unbiased balloons. However, partially different results were revealed: Risk taking gradually increased in the subsequent phase due to adaptation to the changed structure, irrespective of the valence of initial experience. Importantly, the direction of this change in risk-taking behavior contrasts with the behavioral change of the lucky conditions in Experiment 1, and mostly characterizes the second half of the task. The origin of this unexpected finding is discussed in detail in the section entitled as Methodological issues constraining interpretation (see General Discussion).

As participants in the unlucky condition started the subsequent phase with lower pump number due to initially negative experiences, they increased pumping more than other participants. The modulating effect of initial experience was no longer observed in the second half of the task, as the number of pumps became similar across conditions. Results confirmed the expectation following from a previous study ^34^ that the overall effect of negative experience would be more different from baseline than the effect of positive experience. However, the behavioral pattern of the lucky condition was comparable to that of the baseline condition in the subsequent phase. Characteristics of the payoff structure could account for this finding, which is discussed in the General Discussion (see Methodological issues constraining interpretation section). Still, some indication was found in the initial phase that positive experience yielded higher risk taking than baseline experience.

## General Discussion

### Summary of findings

This study examined how initial experience with outcome probabilities influenced later risk-taking behavior in an experience-based risky decision-making task. To this end, we inserted an unsignaled and unexpected change point in the payoff structure of the BART, before and after which outcome probabilities considerably differed. The change point occurred after one-sixth or one-third of trials in the first experiment, and between the initial and subsequent phases in the second experiment. Importantly, the task structure after the change point was unbiased and comparable across experimental conditions, only the relative valence of events differed as a function of previous experience. Although our study was mostly exploratory, several results contradicted the hypotheses based on previous literature. Particularly, both experiments suggested that participants in all conditions adjusted their risk-taking behavior to the changed payoff structure of the task; however, the impact of initial experience was simultaneously observed, especially in the first experiment.

Results of the first experiment showed that initially experiencing negative events led to consistently decreased risk-taking behavior. However, even if initial experience indicated that more cautious behavior was advantageous, risk taking increased after the change in structure. Similarly, if initial experience indicated that high risk could be taken, risk taking decreased after the change. The length of initial experience did not reliably influence the pattern of behavioral change. Results of the second experiment further highlighted that the impact of initial negative experience could be overcome by integrating recent experiences. Participants in all conditions gradually increased their risk taking as the task progressed. Risk-taking behavior after positive initial experience was adjusted to an extent to match behavior after baseline initial experience. This was not the case after negative initial experience: Risk taking remained lower than after baseline initial experience, but it was also adjusted yielding similar behavior across the conditions later in the task.

### Adjustment of risk-taking behavior in the BART

One of our central questions is whether the length of manipulation influences the update of initial beliefs about outcome probabilities. According to the follow-up of the three-way interaction of initial luck, length, and balloon, the length of positive or negative experience does not considerably influence behavior adjustment after manipulation. We found overall higher number of pumps with short than long manipulation only in the bin-wise model (see Supplementary Table S1) and not in the balloon-wise one. Thus, the effect of length is detectable on a larger bot not on a finer time scale. However, this result is not directly related to how risk-taking behavior is adjusted in a changed environment.

Irrespective of length manipulation, results of the first experiment are mostly in line with the findings of Koscielniak, et al. ^28^ and Bonini, et al. ^29^, indicating not only primacy but also recency effects in the BART. In contrast, the study of Yau, et al. ^44^ did not find evidence for change in risk taking when the original BART was followed by a rigged BART with 50% probability of balloon burst. In that study, however, an automatic response version of the task ^54^ was used, which lacks the experience of a dynamic inflation process, and decisions might be more explicitly verbalized.

The adjustment of risk-taking behavior in the BART was found even without changing the payoff structure but providing positive or negative economic forecast (i.e., predictive message that the probability of balloon bursts would decrease or increase in the next task block, see ^30, 34^). Similarly, framing effects (inflate balloons to gain reward or avoid losing reward) without actual structural changes also influenced risk-taking behavior ^36, 55^. By comparing the latter studies with the present one, our contribution is to show that no explicit information or framing is needed to shift individuals’ risk taking to match the changing contingencies of the environment. However, while explicit information allows for abrupt strategy change, experience with the environment gives rise to a smooth adjustment of behavior to the changed contingencies.

Although the present experiments indicate that participants adapt to changed circumstances, initial negative experience still decreases further risk taking. Therefore, many studies using the BART and examining influential factors of risk taking (e.g., the effect of reward type, gender, age, situational factors, and clinical symptoms, see ^35, 38, 39, 56, 57^) could have involuntarily involved the effect of initial experience. If the extent and consistency of initial positive or negative experience were uncontrolled, one could not determine from which phase of the task behavior adjustment should be expected. Thus, if experimental effects and/or group differences were measured relatively early in the task, these could be influenced by initial experience. For instance, as the results of our explorative study presented in the Supplementary Material indicate, even when initial experience is not directly manipulated, individuals would pump the balloons larger during the task if they experienced more successful pumps before balloon bursts within the first five trials (see also Supplementary Fig. S1). Consequently, from a practical angle, it seems that initial experience should be considered when using the BART.

### Some putative cognitive underpinnings of the observed effects

#### Reinforcement learning

The present findings can be interpreted according to the principles of reinforcement learning. It is possible that a shift from goal-directed to habitual behavior occurred, guided by model-based followed by model-free processes ^58, 59^. Individuals might have performed the task according to internal models based on initial experiences about outcome probabilities. By integrating recent experiences, these initial internal models might have gradually changed. During the early phase of task solving with less experience, choices based on a model-based process might be more reliable, since its short-term predictions could be more accurate than long-term predictions of a model-free process. With more experience, the model-based process might become less reliable, since initial internal models could contain inaccuracies that are denoted by unexpected feedback events ^58, 59^. As experience accumulates, a shift is expected from a model-based to a model-free process. The choice behavior found in the first experiment likely reflects such a shift between these processes. This is not that clear in the second experiment, maybe due to weaker (initial) internal models. In addition, based on their robust initial internal models, participants experiencing early negative events could have expected more balloon bursts later in the task, yielding overall more risk-averse behavior in both experiments.

#### Sampling biases

It has been proposed that individuals tend to underweight the probability of rare events when making decisions from experience, such as in the present experiments. Accumulating evidence implies that this tendency originates from relying on relatively small samples and overweighting recent experiences ^6, 13, 31^. Participants in the unlucky conditions sampled the balloons less than those in the lucky conditions, even after the end of manipulation (i.e., lower pump number on balloons that did not burst). Thus, they less likely encountered balloons with very high tolerances, which might have contributed to underestimating the variance of balloon tolerances (i.e., the probability of rare events). Simultaneously, they recently and frequently encountered balloons with “medium” tolerances that were *larger* than those during initial unlucky manipulation. In turn, these frequent events could have been overweighted yielding an increase in risk-taking behavior.

Participants in the lucky conditions with more sampling encountered higher variance of balloon tolerances, which might have eventually yielded more risk-averse behavior ^11^. Similar mechanisms could have contributed to the results of the second experiment, at least in the case of the unlucky condition. However, this theoretical account could not necessarily explain choice behavior of participants in the lucky and baseline conditions in the second experiment, as they sampled the balloons to a larger extent. In their case, local sampling biases could have introduced dynamic changes in risk sensitivity, yielding increased risk taking in time ^12^ (see also the Methodological issues constraining interpretation section below).

#### Memory processes

Sampling biases can be related to memory constraints. In our experiments, participants saw only the accumulated total score and the score collected from the previous balloon on the screen; therefore, no detailed statistics were available about balloon tolerances. Thus, it is conceivable that participants in all conditions did not explicitly track and memorize past events that could have been used to predict the likely outcomes of decisions (but see ^33, 60^). Instead, they changed their strategy in a dynamic and implicit manner, according to experiences on the latest balloons. The weaker memory traces of the initial payoff structure might have contributed to quick adaptation in general, irrespective of the length and valence of early experience (cf. ^20^). The second experiment also indicated the relatively week integration of early memory traces on action-outcome contingencies, as initial manipulation did not considerably alter subsequent risk-taking behavior, at least later in the task.

#### The general anchoring-and-adjustment model

One might attempt to explain our results in terms of the general anchoring-and-adjustment model of belief updating ^8^. In this model, the encoding of evidence (evaluation or estimation), the processing of evidence (step-by-step revisions or the adjustment of the initial anchor based on a series of evidence), and the sensitivity to evidence are critical subprocesses. The combination of these subprocesses together with task characteristics (complexity, length of sampling) determine how the order of information could influence decisions.

Participants could have used both the evaluation (positive or negative deflection from previous experience) and estimation (sequential integration of information) modes of encoding. Both the processing of evidence and the response mode involves step-by-step operations since risky decisions are made at each step of the balloon inflation process. However, it is possible that the end-of-sequence process has also been used (guided by aggregated experience with a series of successful balloon pumps) or a switching between the strategies might have occurred ^19, 39^. The attitudes toward negative and positive evidence might have been constrained by the initial bias. The history of previous balloon pumps and balloon tolerances could be considered as a long series of complex evidence not only in the long manipulation condition but also in the other ones. Considering all processes, the prediction of the anchoring-and-adjustment model would be a force toward primacy effect. Although primacy effect was found in both experiments, recent experience also guided the update of choice behavior, which was not constrained to short initial experience (cf. Table 2 in ^8^). Therefore, the observed findings might not be wholly interpreted along this theoretical model. Also, mapping the structure of the BART onto tasks and concepts that the model has been based on is not obvious.

#### Summary

Taken together, the interpretations of the observed effects remain tentative in terms of the potential cognitive processes. Experimental designs that not only involve the currently investigated factors but also directly manipulate some of the relevant cognitive processes are necessary. In this respect, computational models that account for the effect of initial experience might be considered: With such analyses, initial experience per se could be quantified and other parameters of the task could be compared across participants and groups/conditions while controlling for this effect ^46, 48, 61, 62^.

### Methodological issues constraining interpretation

Two related methodological issues concerning the design of both experiments should be considered: (1) the instability of the remainder/subsequent environment in terms of balloon tolerances (maximum numbers of successful pumps), especially in the second experiment, and (2) the similarity of initial experience between lucky and baseline conditions in the second experiment. These issues are elaborated on below.

The first issue is related to the result showing that the direction of behavioral change differed between the experiments. While lucky experience led to less risk taking after the end of manipulation in the first experiment, it led to more risk taking in the second experiment. This between-experiments difference could be explained by the different balloon tolerances in the remainder and subsequent phases of each experiment. In the second experiment, the variability of mean balloon tolerances across individual bins was considerable (see Table 2 and Supplementary Fig. S3). Particularly, mean balloon tolerances were relatively high in Bin3 (12.2), Bin4 (13.2), and Bin5 (11.8), intermediate in Bin1 (8.4) and Bin6 (9.2), and low in Bin2 (7).

Therefore, regarding the association between experimental conditions and mean balloon tolerances, the second half of the subsequent experience resembled to the lucky initial experience. Meanwhile, bins were more consistent (except for Bin2) throughout the whole task in the case of baseline experience. Mean tolerances of the first two bins were higher than that of unlucky initial experience but lower than that of lucky initial experience. Still, these bins could have been “lucky” or at least “neutral” experience for participants of the unlucky condition and “unlucky” experience for participants of the lucky condition. Altogether, the inconsistent or instable nature of the subsequent experience could have altered how participants adapted to the changed environment.

Particularly, the change of behavioral measures also reflected the change of balloon tolerances: Balloon pumps were increased, and the frequency of balloon bursts was decreased in Bin3, Bin4, and Bin5 in all conditions of the second experiment. Meanwhile, decreased balloon pumps were observed in Bin2 relative to Bin1 in the case of lucky and baseline experiences (Fig. 2b, Supplementary Fig. S3). Thus, the bin-by-bin variation of mean tolerance values could have determined adaptation: The positive change in risk-taking behavior was probably caused by the positive change in the payoff structure in the second half of the task. Indeed, after initial experience, risk taking became advantageous for all conditions in the subsequent environment, at least from Bin3. The more cautious (lucky) or relatively unchanged (baseline, unlucky) risk-taking behavior in the first two bins can also be explained by this reasoning.

In contrast to the second experiment, mean balloon tolerances in the remainder of the task were slightly lower and less variable in the first experiment (first experiment: *M* = 9.07, *SD* = 1.05; second experiment: *M* = 10.30, *SD* = 2.45, see also Table 2 and Supplementary Fig. S2). These balloon tolerances were intentionally sampled from a narrower uniform distribution to obtain “medium” balloon sizes, which could yield lower variability. More importantly, these values in the first experiment were considerably lower (higher) than that of the initial lucky (unlucky) balloons. Therefore, similarly to the second experiment, the change in the payoff structure could account for the decreased (lucky) and increased (unlucky) risk-taking behavior in the first experiment.

As tolerance values were identical across participants in both experiments, the described variability could result in a systematic confound that influences the interpretability of the observed effects, at least in the second experiment. Therefore, to test the original research questions more rigorously, additional carefully designed experiments should be performed. A proper experimental design warrants consistent, stable experiences across conditions and time bins in the novel environment that follows manipulation. In the present study, however, it seems that behavior adjustment depended on the change in tolerance values irrespective of the type of initial experience, which might also be regarded as indicative of how primacy effect can be overcome by the integration of recent experiences.

The second methodological issue is related to the result showing that the behavioral change was similar between the lucky and baseline conditions and differed from the unlucky condition in the second experiment. This suggests that the processing of negative events in the unlucky condition differently influenced later risk-taking behavior as compared with the other conditions. On the one hand, this asymmetry was expected since stronger effect of negative manipulation has already been found in previous BART-studies ^29, 30, 34^. On the other hand, it is probable that the origin of this asymmetry also lies in how balloon tolerances were selected.

While the maximum number of successful pumps was not significantly higher in the lucky than in the baseline condition in the initial phase, it was much lower in the unlucky condition (see Stimuli, design, and procedure section). Therefore, initial experience could have been similar between lucky and baseline conditions, yielding comparable risk-taking behavior in the subsequent phase of the task. The above-described change in the payoff structure of the subsequent phase could have also played a role in this asymmetry. At the same time, it should be noted that both the balloon-wise and bin-wise model result showed that pump number was increased by later balloons of the initial phase in the lucky condition as compared with the baseline condition (see also Fig. 2a).

Beyond methodological issues that could account for the change in risk-taking behavior in the second experiment, the effect of weak memory traces is also a possible explanation (see the earlier section on memory processes). Separating task phases could have unintentionally suggested that novel circumstances would apply after the short pause. Thus, after some (probably surprising) experience in the early trials of the subsequent phase, participants of all conditions might have perceived this phase as an entirely independent task and increased pump number as usually observed in the original BART ^2, 4, 35, 49, 63–65^. Nevertheless, in further experiments, clearly distinguishable initial experiences should be applied.

### Limitations

The design of the experiments inherently limited what participants could initially learn about the balloons. Participants’ baseline belief about the pump number that balloons could have tolerated was not fixed a priori before the manipulation. They might have established this belief based on the experiences gathered during the first few trials (cf. ^12, 66^). However, it was limited to what size they could pump the balloons at the beginning of the task; and it was also unknown to what degree they were prone to pump these balloons.

Relatedly, an inherent limitation of the BART is that it provides one-sided censored data, since burst events terminate trials perforce ^4^. Therefore, burst trials cannot reveal to what extent participants were willing to take risks or what were the underlying processes that determined the pump number on those trials (e.g., risk taking, impulsivity, task engagement or motivation, and reduced sensitivity to negative feedback). Moreover, by excluding these trials, the reliability of data could slightly decrease (i.e., fewer trials), and this remaining data would still include any effects that the burst events had on subsequent trials ^46^.

Participants received neither compensation for participation nor bonus based on task performance because of practical reasons. Recent work has implied that not only the behavioral indices but also the neurophysiological correlates of negative feedback processing are enhanced if real money instead of hypothetical reward is used in the BART ^57, 67^. Nevertheless, according to the impressions of the experimenters during debriefing, participants were still motivated to perform the task as if gains and losses were real.

### Conclusions

This study confirmed that the combination of both primacy and recency effects contributes to experience-based risky decision making. Moreover, the study revealed that extended early experience does not induce more profound primacy effect than brief early experience. Instead, choice behavior is adjusted by integrating recent experiences of outcome probabilities into the internal models of the decision environment. Therefore, while choice behavior remains risk averse after early negative events, this attenuates with the accumulation of new experiences. Several factors could underlie the observed effects, such as a shift from goal-directed to habitual behavior, the overweighting of recent experience, and weak memory traces on initial outcome probabilities; all of which necessitate further investigation. In sum, by showing that initial beliefs are updated to promote adaptation, the present results extend the understanding of uncertain decision making in volatile environments.

## Data availability

All data and analyses codes to reproduce the results are available in the following online repository: https://osf.io/bwzfa/?view_only=f0920ce7759144ceb4be81318cc76f75

## Acknowledgements

This research was supported by the National Brain Research Program (project 2017-1.2.1-NKP-2017-00002, PI: D. N.); the Hungarian Scientific Research Fund (OTKA FK 124412, PI: A. K., OTKA PD 124148, PI: K. J., OTKA K 128016, PI: D. N.); the IDEXLYON Fellowship of the University of Lyon as part of the Programme Investissements d’Avenir (ANR-16-IDEX-0005 to D. N.); the János Bolyai Research Scholarship of the Hungarian Academy of Sciences (to A. K. and K. J.); and the Postdoctoral Fellowship of the Hungarian Academy of Sciences (to A. K.). We thank Vera Varga for providing helpful suggestions on data analysis and the reviewers for their illuminating comments.

## Author contributions

A. K. designed the study, created the scripts running the experiments, analyzed data, and wrote the manuscript; Z. K. designed the study, analyzed data, contributed to the interpretation of the results, wrote the manuscript, and critically revised previous versions of the manuscript; Á. T. designed the study, supervised data acquisition, analyzed data, wrote the manuscript, and critically revised previous versions of the manuscript; N. É. designed the study, analyzed data of the explorative study presented in the Supplementary Material, contributed to the interpretation of the results, and critically revised previous versions of the manuscript; K. J. designed the study and critically revised previous versions of the manuscript, E. T-F. analyzed data, rated the interview questions, and revised previous versions of the manuscript; V. Cs. revised previous versions of the manuscript and provided intellectual support; D. N. designed the study, critically revised previous versions of the manuscript, and provided intellectual and financial support. All authors read and approved the final version of the manuscript.

