## Supplementary Material for "Adaptation to recent outcomes attenuates the lasting effect of initial experience on risky decisions"

**Explorative study on the effect of initial experience**

For explorative purposes, we tested the effect of initial experience on risk-taking behavior in a large sample of young adults. In this earlier unpublished study of our lab, the BART was used without experimentally controlling initial experience on action-outcome probabilities.

**Method**

**Participants.** This study involved 180 healthy young adults, who were undergraduate students of Eötvös Loránd University, Budapest, Hungary. Data of 156 participants (see Table 1 in the main text) were analyzed regarding the initial experience (for details, see Data Analysis section below). All participants had normal or corrected-to-normal vision and none of them reported a history of any neurological and/or psychiatric condition. They provided written informed consent before enrollment. The study was approved by the United Ethical Review Committee for Research in Psychology (EPKEB) in Hungary and by the research ethics committee of Eötvös Loránd University, Budapest, Hungary. It was conducted in accordance with the Declaration of Helsinki. Participants received course credits for their participation.

**Stimuli, design, and procedure.** Participants had to inflate 30 balloons in this version of the BART. The structure and appearance of the task was the same as described in previous studies ^1-5^; and it was written in E-Prime 2.0 (Psychology Software Tools, Inc.). As in Experiments 1–2 presented in the main text, participants were asked to achieve as high score as possible by inflating empty virtual balloons on the screen. In this task version, however, each successful pump increased the potential reward but also the probability to lose the accumulated score because of balloon burst. The maximum number of successful pumps for each balloon were not fixed. Instead, although the regularity determining balloon bursts was unknown to participants, it followed three principles: (1) balloon bursts for the first and second pumps were disabled; (2) the maximum number of successful pumps for each balloon was 19; (3) the probability of a balloon burst was 1/18 for the third pump, 1/17 for the fourth pump, and so on for each further pump until the 20th, where the probability of a balloon burst was 1/1.

The BART was administered as part of a larger project including neuropsychological tests and questionnaires measuring the different aspects of learning, cognition, personality, and social behavior ^6-8^. This project included three experimental sessions; participants performed the BART during the middle session. Here we only report results of the BART.

**Data analysis.** To quantify initial experience, we determined the number of balloon bursts within the first five balloons. Then, we determined the mean number of pumps at which these bursts occurred. After that, we calculated a Pearson’s linear correlation between the mean number of pumps until burst within the first five balloons and the mean adjusted number of pumps across all balloons (MAP; mean number of pumps on balloons that did not burst). This analysis involved only 156 participants of the 180; those, who experienced at least one balloon burst within the first five balloons.

**Results and Discussion**

The mean number of pumps until burst within the first five balloons significantly correlated with the MAP, *r*(154) = .503, *p* < .001, suggesting that initial luck was associated with risk-taking behavior on the BART (see Fig. S1). The later participants experienced balloon burst initially, the more they tended to explore the task’s structure and take risk. Thus, the first five experienced outcomes influenced participants’ risky choices. However, in this analysis, the designation of the exact end of initial experience at five balloons was arbitrary and only served explorative purpose. In addition, the MAP was calculated for all the 30 balloons (see Table 1 in the main text).

**Supplementary Tables**

**Table S1.** Summary of the bin-wise version of Model 5 related to Experiment 1.

| *Fixed effects* | Estimate | *SE* | df | *t*-value | *p*-value |
| --- | --- | --- | --- | --- | --- |
| (Intercept) | 8.77 | 0.31 | 134.57 | 27.93 | < 0.001 |
| **Luck** | **-4.26** | **0.50** | **214.11** | **-8.49** | **< 0.001** |
| **Length** | **1.22** | **0.45** | **138.08** | **2.73** | **0.007** |
| **Bin 2** | **1.71** | **0.28** | **1396.88** | **6.13** | **< 0.001** |
| Bin 3 | -0.16 | 0.32 | 1399.58 | -0.48 | 0.631 |
| **Bin 4** | **-1.03** | **0.29** | **1403.95** | **-3.62** | **< 0.001** |
| Bin 5 | -0.61 | 0.35 | 1403.96 | -1.73 | 0.084 |
| **Bin 6** | **-1.39** | **0.30** | **1399.50** | **-4.70** | **< 0.001** |
| Luck * Length | -0.99 | 0.70 | 205.99 | -1.41 | 0.159 |
| **Luck * Bin 2** | **-1.79** | **0.47** | **1396.96** | **-3.81** | **< 0.001** |
| Luck * Bin 3 | 0.76 | 0.48 | 1398.19 | 1.58 | 0.114 |
| **Luck * Bin 4** | **3.39** | **0.45** | **1400.61** | **7.49** | **< 0.001** |
| **Luck * Bin 5** | **2.69** | **0.53** | **1400.42** | **5.11** | **< 0.001** |
| **Luck * Bin 6** | **2.88** | **0.47** | **1397.88** | **6.17** | **< 0.001** |
| **Length * Bin 2** | **-2.11** | **0.44** | **1398.10** | **-4.78** | **< 0.001** |
| Length * Bin 3 | -0.45 | 0.44 | 1397.78 | -1.04 | 0.299 |
| Length * Bin 4 | -0.15 | 0.48 | 1400.72 | -0.31 | 0.760 |
| Length * Bin 5 | -0.61 | 0.48 | 1400.12 | -1.29 | 0.198 |
| Length * Bin 6 | -0.34 | 0.43 | 1398.10 | -0.78 | 0.436 |
| **Luck * Length * Bin 2** | **3.54** | **0.67** | **1396.93** | **5.28** | **< 0.001** |
| **Luck * Length * Bin 3** | **2.20** | **0.65** | **1397.17** | **3.38** | **0.001** |
| Luck * Length * Bin 4 | 0.27 | 0.71 | 1398.89 | 0.39 | 0.701 |
| Luck * Length * Bin 5 | 0.45 | 0.71 | 1398.48 | 0.64 | 0.525 |
| Luck * Length * Bin 6 | 0.86 | 0.66 | 1398.02 | 1.29 | 0.197 |
| *Random effects* | Variance | *SD* |  |  |  |
| [Participants](Intercept) | 1.29 | 1.14 |  |  |  |

*Note*. The dependent variable was the number of pumps on each balloon that did not burst. This model was fit to all balloons of all conditions. For a more detailed description of the model, see main text. With the treatment contrast, the reference level of a given factor was used to estimate the other levels. These were lucky, long, and Bin 1. Significant effects are in **bold**. *SE*: standard error; *SD*: standard deviation.

**Supplementary Figures**

MAP

| Number of pumps until burst within the first five balloons | MAP |
| --- | --- |
| 5 | 7.61 |
| 4 | 8.29 |
| 7 | 8.06 |
| 5 | 9.68 |
| 7 | 10.8 |
| 4 | 8.75 |
| 14 | 12.13 |
| 7 | 7.43 |
| 6 | 6.69 |
| 8 | 9.27 |
| 5 | 6.05 |
| 6 | 6 |
| 4 | 7.94 |
| 4 | 5.94 |
| 5 | 7.62 |
| 9 | 8.08 |
| 6 | 6.85 |
| 5 | 7.39 |
| 13 | 11.4 |
| 4 | 4.29 |
| 5 | 7.53 |
| 13 | 7 |
| 10 | 7.18 |
| 7 | 6 |
| 3.67 | 9 |
| 6 | 8.33 |
| 13.67 | 8.69 |
| 6 | 8.38 |
| 8 | 6.5 |
| 10 | 11.61 |
| 6 | 11.43 |
| 3 | 3.37 |
| 12 | 10.6 |
| 10.33 | 7.62 |
| 8 | 11.39 |
| 5 | 6.95 |
| 8.5 | 9.8 |
| 3.5 | 9.12 |
| 9 | 8.15 |
| 6.5 | 7.59 |
| 10.25 | 8.89 |
| 3 | 7.19 |
| 4 | 6.1 |
| 3.5 | 2.96 |
| 7 | 6.57 |
| 11.67 | 10.69 |
| 4 | 9.42 |
| 4.5 | 9.15 |
| 5 | 6.94 |
| 6 | 4.91 |
| 4 | 8.94 |
| 5.5 | 10.05 |
| 6.25 | 11.3 |
| 6 | 11.17 |
| 8.67 | 11.09 |
| 9 | 12.92 |
| 13 | 13.5 |
| 7 | 8.36 |
| 5 | 5.86 |
| 12 | 9.07 |
| 4 | 6.86 |
| 10 | 7.68 |
| 4.5 | 5.9 |
| 5 | 8.32 |
| 9.67 | 9.56 |
| 7 | 8.72 |
| 7 | 10.37 |
| 5 | 6.3 |
| 3 | 5.08 |
| 6 | 6.71 |
| 6 | 6.94 |
| 4 | 5.04 |
| 7.5 | 6.57 |
| 3.5 | 5.9 |
| 3.5 | 9.53 |
| 7 | 7.15 |
| 3 | 3.5 |
| 8 | 7.5 |
| 6 | 8.8 |
| 9.5 | 10.6 |
| 4 | 9.63 |
| 6.5 | 9.14 |
| 6 | 9.06 |
| 12.5 | 7.06 |
| 13.5 | 7.28 |
| 4 | 5.7 |
| 4 | 4.88 |
| 6.5 | 5.29 |
| 12.33 | 7.61 |
| 12 | 13.71 |
| 12 | 14.67 |
| 5.5 | 11.09 |
| 10.5 | 10.39 |
| 8 | 9.53 |
| 5.33 | 6.77 |
| 9.33 | 8.4 |
| 3 | 6.22 |
| 4.5 | 5.33 |
| 8 | 10 |
| 7 | 5.26 |
| 6.5 | 9.29 |
| 5 | 7.92 |
| 4 | 5.25 |
| 7 | 8.9 |
| 8.5 | 11.67 |
| 5 | 7.64 |
| 6 | 8.58 |
| 5 | 10.93 |
| 4 | 6.25 |
| 11 | 10.82 |
| 7.67 | 7.67 |
| 5.33 | 8.71 |
| 7 | 11.44 |
| 4 | 10.7 |
| 5 | 10.17 |
| 3 | 11.25 |
| 4 | 7.62 |
| 5 | 7.3 |
| 3 | 6.14 |
| 13.67 | 10.93 |
| 4 | 7.48 |
| 5.5 | 7.89 |
| 4 | 4.5 |
| 6 | 7.95 |
| 3 | 6 |
| 5 | 6.62 |
| 5 | 6.22 |
| 6 | 6.89 |
| 7 | 8.05 |
| 4 | 7.32 |
| 4 | 5.05 |
| 7.33 | 7.44 |
| 3 | 3.76 |
| 9 | 7.48 |
| 9.33 | 11.67 |
| 7.33 | 9.6 |
| 10 | 12.36 |
| 10.33 | 10.38 |
| 4 | 7.72 |
| 7.67 | 5.95 |
| 7.67 | 7.63 |
| 7.33 | 4.25 |
| 15 | 8.94 |
| 8 | 6.79 |
| 8 | 9 |
| 11 | 8.12 |
| 10 | 9.17 |
| 5.33 | 12.83 |
| 10.25 | 11.5 |
| 6 | 11.47 |
| 7 | 9.14 |
| 10 | 10.92 |
| 11 | 12.36 |
| 3 | 6.45 |
| 6 | 5.94 |
| 15 | 11.42 |

**Figure S1.** Scatter plot illustrating the association between the mean number of pumps until balloon burst within the first five balloons and mean adjusted number of pumps (MAP) in an independent study with a large sample of young adults.

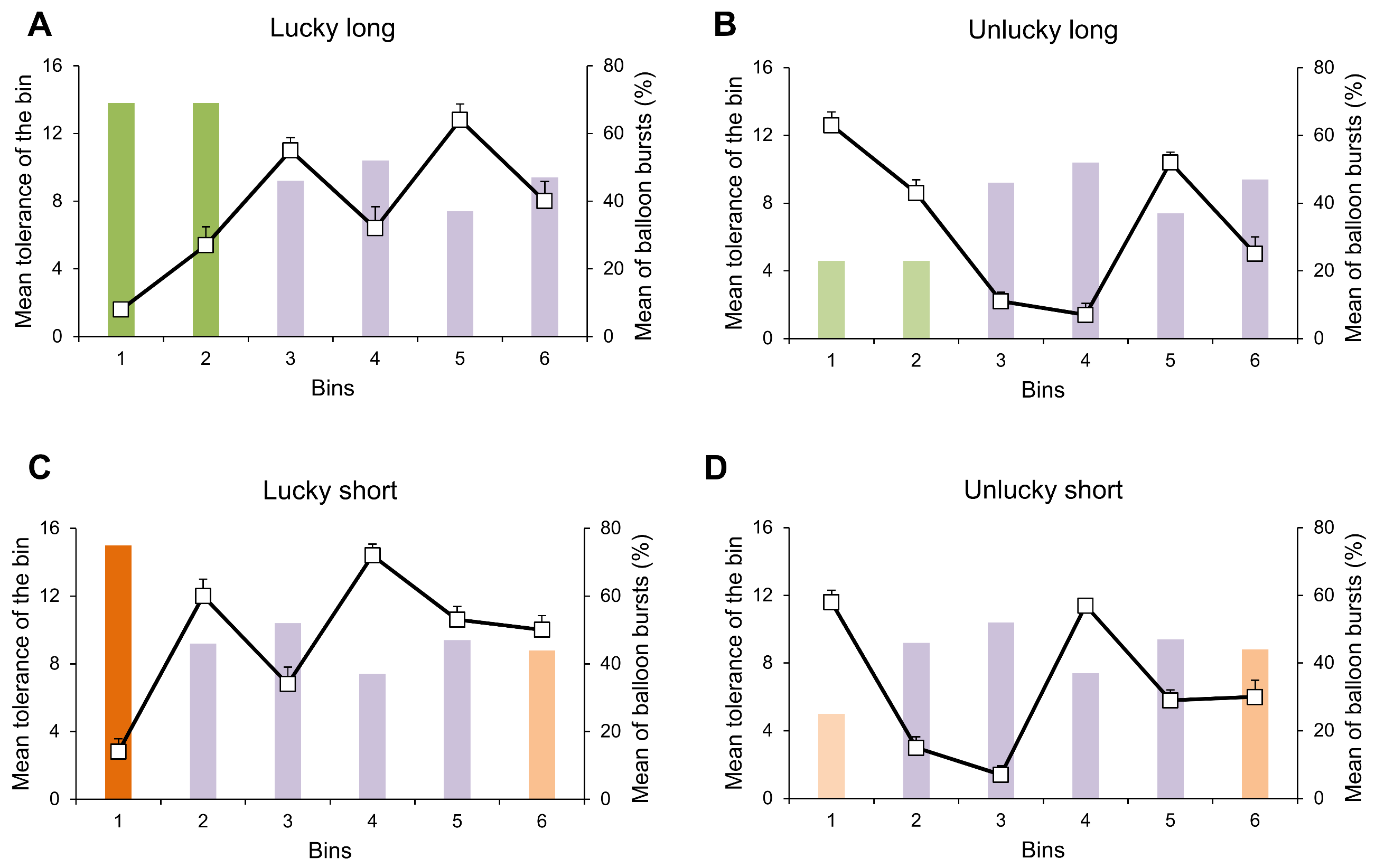

**Figure S2. Balloon tolerances and balloon bursts in Experiment 1.** Bar charts and left y-axes represent mean balloon tolerances in each time bin (5 consecutive balloons). Line charts with white square symbols and right y-axes represent the mean percentages of balloon bursts in each time bin (i.e., if all five balloons burst, this was 100%). These two indices of the experiment are plotted against one another in each condition (A: lucky long, B: unlucky long, C: lucky short, D: unlucky short). Time bins with the same mean balloon tolerances across conditions are violet-colored. The first two bins of the long manipulation are dark green (A, lucky long) and light green (B, lucky short), respectively. The first bis of the short manipulation are dark orange (C, lucky short) and light orange (D, unlucky short), respectively. The different colors indicate that these initial bins slightly differ in mean tolerances between the long and short manipulation conditions. The final bins of the lucky short and unlucky short conditions have the same mean tolerance that is different from the other values. Error bars denote standard error of mean. (The visibility of error bars could be poor because of their low value and the resolution of the scale.) Note that when balloons tolerated a higher number of pumps in a given time bin, the frequency of balloon bursts was higher in the next time bin, even though the actual mean tolerance of this bin was lower. Then, risk taking was calibrated to the recent outcomes yielding a reduced frequency of balloon bursts in the next bin. This pattern was pronounced in the lucky conditions, but it was also apparent in the unlucky conditions after participants had overcome initial experience.

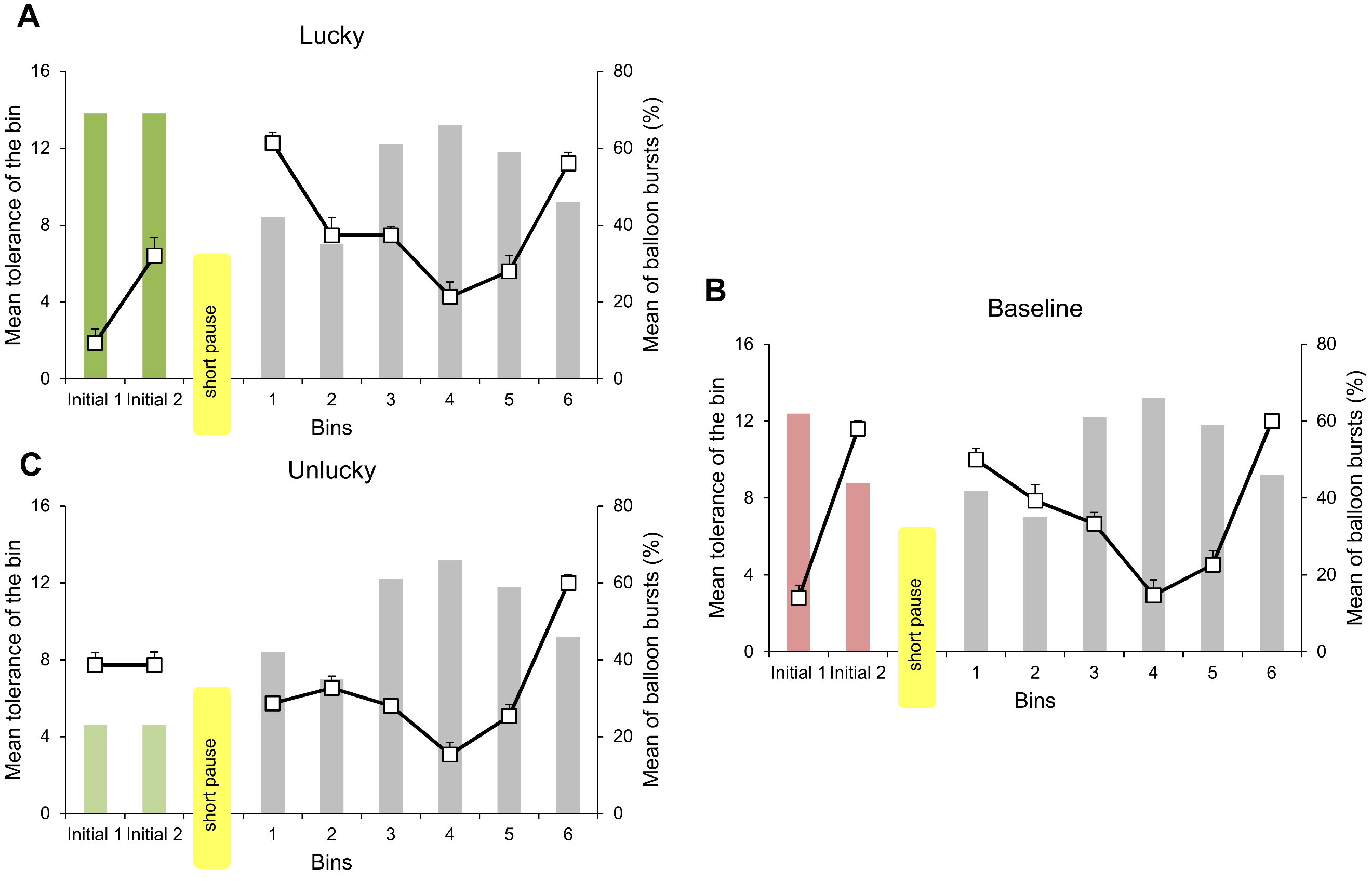

**Figure S3.** **Balloon tolerances and balloon bursts in Experiment 2.** Bar charts and left y-axes represent mean balloon tolerances in each time bin (5 consecutive balloons). Line charts with white square symbols and right y-axes represent the mean percentages of balloon bursts in each time bin (i.e., if all five balloons burst, this was 100%). These two indices of the experiment are plotted against one another in each condition (A: lucky, B: baseline, C: unlucky). Vertical yellow areas separate the initial and subsequent phases of the task, each containing two and six time bins, respectively. This short pause lasted a few seconds until the experimenter initiated the next phase of the task. Time bins with the same balloon tolerances across conditions are grey-colored. The first two bins of the lucky condition (A) are dark green as they were the same as in the lucky long condition of Experiment 1 (see Figure S2A). The first two bins of the unlucky condition (C) are light green as they were the same as in the unlucky long condition of Experiment 1 (see Figure S2A). These were different from the tolerance values of the first two bins of the baseline condition (B). Error bars denote standard error of mean. (The visibility of error bars could be poor because of their low value and the resolution of the scale.) Note that the bin-by-bin adaptation was not as emphasized in Experiment 2 as in Experiment 1, possibly due to the between-experiments differences in balloon tolerances.
